# Tissue-Specific Co-Expression Patterns of BAF Complexes Provide Regulatory Insights Across Human Tissues with Implications for Endocrine and Non-Endocrine Functions

**DOI:** 10.1101/2025.07.07.663564

**Authors:** Xiaowei Dong, Neshatul Haque, Jessica B Wagenknecht, Michael T Zimmermann

## Abstract

BRG1/BRM-associated factor (BAF) chromatin remodeling complexes are essential for normal endocrine function and are implicated in various metabolic and developmental disorders. However, the full range of chromatin-based regulatory modules in endocrine development remains unclear. We developed a computational pipeline to analyze bulk RNA-seq data from 54 human tissues and constructed tissue-specific co-expression networks for 30 core BAF complex genes. Weighted gene co-expression network analysis (WGCNA) and Louvain clustering identified gene modules for each tissue, which we compared to 46 curated BAF subcomplex gene sets using Jaccard similarity.

In metabolically active non-endocrine tissues (kidney, skeletal muscle, vasculature, fibroblasts), we observed strong co-expression with canonical BAF (cBAF) and polybromo-associated BAF (pBAF) modules. Central nervous system tissues were dominated by neuron-specific BAF (nBAF) modules. Endocrine tissues (e.g., thyroid, adrenal) and gastrointestinal epithelia displayed co-expression profiles resembling smooth muscle–like BAF and pBAF modules, suggesting chromatin programs that integrate hormone secretion with contractile and barrier functions. These patterns show that each tissue exhibits a distinct, non-random combination of BAF subcomplexes, potentially reflecting its functional chromatin state.

Our results demonstrate that tissue-specific gene expression profiling can reveal differences in protein complex regulation. The modular deployment of BAF chromatin remodeling complexes appears tailored to the functional demands of each organ. This study lays a foundation for further investigation of epigenetic regulation in endocrine development and disease and provides a framework for identifying tissue-specific chromatin remodeling strategies.

**Plain Language Summary (optional):** The US National Institutes of Health have invested in large-scale measurements of how different human body tissues use their genetic material. This study pilots use of these data to understand better how various genes come together to form nanomachines that in turn regulate the same genetic material.

## Introduction

Epigenetic regulatory complexes are central to the development and functional maintenance of all body tissues. These complexes orchestrate tissue-specific gene expression patterns that drive the differentiation of endocrine cells and ensure the precise synthesis, release, and responsiveness to hormones throughout life. For example, persistent or aberrant modification to the chromatin remodeling complex BAF (BRG1/BRM-associated factor), can disrupt endocrine homeostasis and contribute to metabolic and developmental disorders. Moreover, epigenetic regulation integrates environmental cues and metabolic states with endocrine function, mediating phenotypic plasticity and enabling adaptive responses to physiological and environmental changes. Thus, epigenetic regulatory complexes are not only foundational for establishing endocrine tissue identity but also for dynamically modulating hormone action and maintaining systemic metabolic balance.

Understanding gene expression regulation is fundamental to deciphering how cells control their structure, function, and response to cues. Current knowledge remains incomplete due to the immense complexity of regulatory networks and the diversity of mechanisms involved. Large-scale sequencing studies, such as using transcriptomics to measure the levels of all messenger RNAs, now provide unprecedented opportunities to systematically map gene co-expression patterns across diverse conditions and cell types. These patterns are valuable because genes that are co-expressed often participate in shared regulatory programs, functional pathways, or are controlled by common transcription factors or epigenetic mechanisms. In brief, co-expression can indicate co-regulation. The justification for using co-expression as a proxy for co-regulation lies in the principle that coordinated expression across multiple samples or time points frequently reflects underlying regulatory relationships, such as shared cis-regulatory elements, chromatin states, or trans-acting factors, thereby enabling researchers to infer functional gene modules and predict regulatory interactions from high-dimensional data.

BAF complexes are fundamental to chromatin remodeling, playing a crucial role in regulating gene expression by modifying chromatin accessibility. BAF exists as specialized versions. For example, the canonical (cBAF), non-canonical (ncBAF), and poly-bromo containing (pBAF) versions are each tailored for distinct regulatory functions. Their diverse subunit compositions enable dynamic interactions with transcriptional machinery, influencing key biological processes such as cell differentiation, development, and disease pathogenesis [1]. Also, there is a typical order of assembly with branch points that decide between the three versions [2].

Amazingly, the composition of BAF complexes varies significantly between cell types, reflecting the unique transcriptional landscapes required for cellular identity. Subunits such as BAF45A (DPF1), BAF45B (DPF2), BAF45C (DPF3), BAF45D (PHF10), BAF60A (SMARCD1), BAF60B (SMARCD2), BAF60C (SMARCD3), and the BCL7 family (BCL7A, BCL7B, BCL7C), exhibit distinct expression patterns, aligning with the specific needs of each cell type. This variability highlights the cell-type-specific regulation of chromatin architecture, as evidenced by large-scale epigenomic studies such as the Roadmap Epigenomics Project [3]. The differential expression of BAF subunits is particularly critical in tissue-specific functions, especially within the endocrine system, where precise gene regulation is essential for hormone synthesis and secretion. We hypothesize that BAF complex versions are adapted in composition to meet the transcriptional demands of endocrine tissues. For example, from other tissues, BAF60C plays a key role in cardiac development, facilitating the recruitment of heart-specific enhancers during early embryogenesis [4]. Beyond normal development, disruptions in BAF complex function are implicated in diseases such as Coffin-Siris syndrome, where mutations in BAF components lead to developmental abnormalities, including endocrine dysfunctions like hypothyroidism or growth hormone deficiency [5]. These findings underscore the critical role of BAF complexes in maintaining tissue homeostasis, demonstrating their broader significance in both normal physiology and disease pathology.

Since the introduction of transcriptome-wide profiling, module detection methods have become essential for analyzing large-scale gene expression data. Modules refer to groups of genes with similar expression patterns, often reflecting functional relationships and co-regulation. Modules have an advantage over co-expression because the group-wise pattern is typically more biologically relevant than individual pairs of genes. They help researchers understand gene interactions, infer regulatory relationships between transcription factors and target genes [6–9], and enhance genome annotation through the guilt-by-association principle [6,9]. Additionally, module detection provides insights into disease mechanisms and progression [10,11]. To identify gene modules, various unsupervised machine learning approaches have been developed. Clustering, which groups genes based on similarity in expression profiles, remains one of the most widely used methods [6,10]. However, alternative approaches provide more refined insights. Decomposition techniques, such as Principal Component Analysis (PCA) and Independent Component Analysis (ICA), extract latent regulatory patterns by breaking down expression data into discrete group-wise components. Meanwhile, Network-based approaches like gene co-expression networks (GCNs), where nodes represent genes and edges encode the strength of their pairwise co-expression, provide a systems-level view of transcriptional regulation [12].By partitioning networks into modules, defined as groups of genes with relatively high intra-group weights compared to inter-group weights, one can reduce high-dimensional transcriptomic data to functionally coherent units [10].

In this study, we used the WGCNA framework to build a signed and weighted co-expression matrix among 30 BAF complex genes and applied the Louvain algorithm to identify non-overlapping modules by maximizing the Newman–Girvan modularity score [13,14]. This combination of WGCNA’s robust, weighted network construction with Louvain’s rapid, scalable community detection allows sensitive recovery of biologically relevant gene modules from large, heterogeneous transcriptomic datasets. The current pilot study aims to uncover the regulatory logic underlying normal endocrine tissue development and function, as well as to highlight disruptions associated with heritable and malignant endocrine diseases. We find that BAF complex genes group into clear, tissue-specific modules that match known chromatin remodeling types, such as canonical, neuron-specific, smooth muscle–like, and non-canonical forms. Ultimately, this approach promises to reveal novel data-driven insights into the more detailed and nuanced molecular mechanisms driving endocrine health and disease, and to identify candidate regulatory modules for further functional investigation.

## Materials and methods

### Data acquisition and preprocessing

The GTEx bulk RNA-seq dataset (V10 release) was obtained from the GTEx portal [15]. The dataset includes 59,033 genes from 19,618 tissue-specific samples. Reliable co-expression estimates require sufficient sample sizes. Smaller than100 samples yields unstable correlation estimates and poor soft-threshold fits. Tissues with fewer than 100 RNA-seq samples were removed (Cervix, Ectocervix, n = 24; Cervix, Endocervix, n = 23; Kidney, Medulla, n = 11; Bladder, n = 77; Fallopian Tube, n = 29). The remaining 49 tissues provided 18,452 samples for analysis. To produce a high-quality input for network analysis, we applied a multi-step preprocessing pipeline. First, we filtered the expression matrix to retain only the 30 BAF complex protein-coding genes (ACTB, ACTL6A, ACTL6B, ARID1A, ARID1B, ARID2, BCL7A, BCL7B, BCL7C, BICRA, BICRAL, BRD7, BRD9, DPF1, DPF2, DPF3, PBRM1, PHF10, RB1, SMARCA2, SMARCA4, SMARCB1, SMARCC1, SMARCC2, SMARCD1, SMARCD2, SMARCD3, SMARCE1, SS18, SS18L1), using HGNC gene symbols.

We generated this list of genes by the union of literature [16,17] and BAF entries from the epiFactors database [18] and CORUM [19]. Next, raw TPM values were transformed as log_2_ (TPM + 1) to dampen heteroscedasticity. Density histogram plots before and after transformation are shown in Additional Figures **S1–S2**.To minimize technical bias introduced by sequencing center, ComBat, which implements an empirical-Bayes framework[20], was applied to adjust for batch effects stemming from different sequencing centers, using nucleic acid isolation batch as the normalization batch factor and including RNA Integrity Number and total ischemic time (minutes from donor ischemia to stabilization) as covariates to preserve biological variation (Additional Figures **S3–S4**). Thus, our gene expression analysis approach is highly robust and focused on BAF as a proof-of-concept for datamining differences in protein complexes regulation across human tissues. A schematic overview of these steps is provided in **Figure 1**.

**Figure 1.**
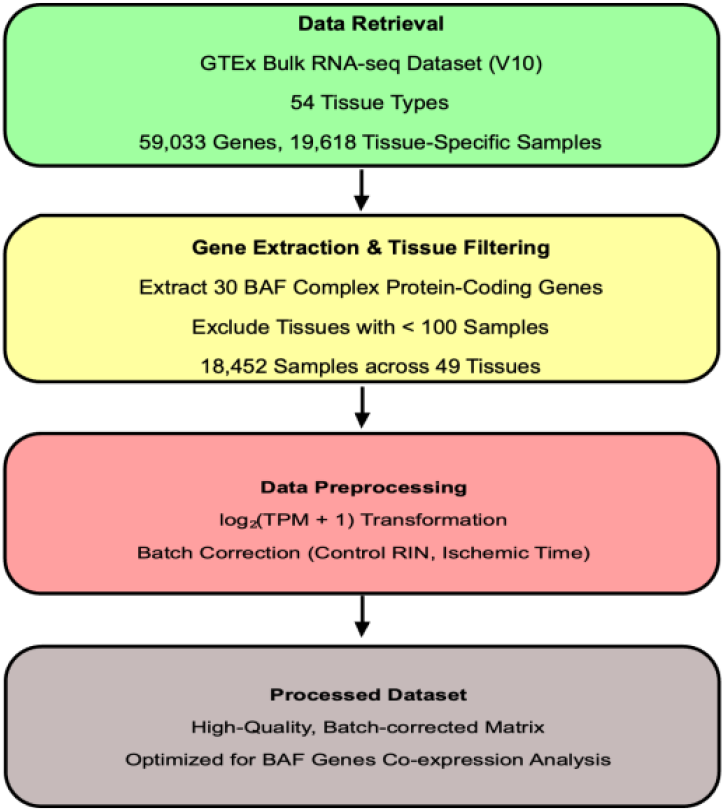
Overview of data preprocessing workflow. Raw GTEx V10 bulk RNA-seq data (19,618 samples, 54 tissues, 59,033 genes) was filtered to include only the 30 BAF complex genes and tissues with at least 100 samples (18,452 samples, 49 tissues remaining). The data were then log_2_(TPM + 1)-transformed and batch-corrected, adjusting for RNA Integrity Number and total ischemic time (minutes from donor ischemia to stabilization), to yield a high-quality, BAF-focused co-expression matrix optimized for downstream analysis

### Endocrine vs Non-Endocrine Annotation

Human tissues were categorized into one of four endocrine classes (primary, secondary, supporting, or other). The first group includes major hormone-producing organs like the adrenal glands, pancreas, pituitary, thyroid, and reproductive organs (testis and ovary)—these are the body’s main hormone factories [21]. The second group covers organs that make hormones but also do other important jobs, such as the liver, kidneys, heart, fat tissue, and placenta [22]. The third group consists of tissues that help control hormone activity without being major producers themselves. This includes the hypothalamus, pineal gland, and parts of the digestive system like the small intestine, large intestine, and stomach [23]. Everything else, such as most brain areas (except the hypothalamus and pineal gland), muscles, lungs, esophagus, and colon, went into the fourth group since they don’t play major roles in making or controlling hormones [24].

We used this grouping system when analyzing how BAF complex genes are expressed differently between hormone-related and non-hormone-related tissues.

### BAF genes co-expression network analysis

We implemented a multi-stage computational pipeline to identify robust functional modules within the BAF complex. An overview of Co-Expression Network Construction, Module Detection, and Community Validation and Visualization is presented in **Figure 2.**

**Figure 2.**
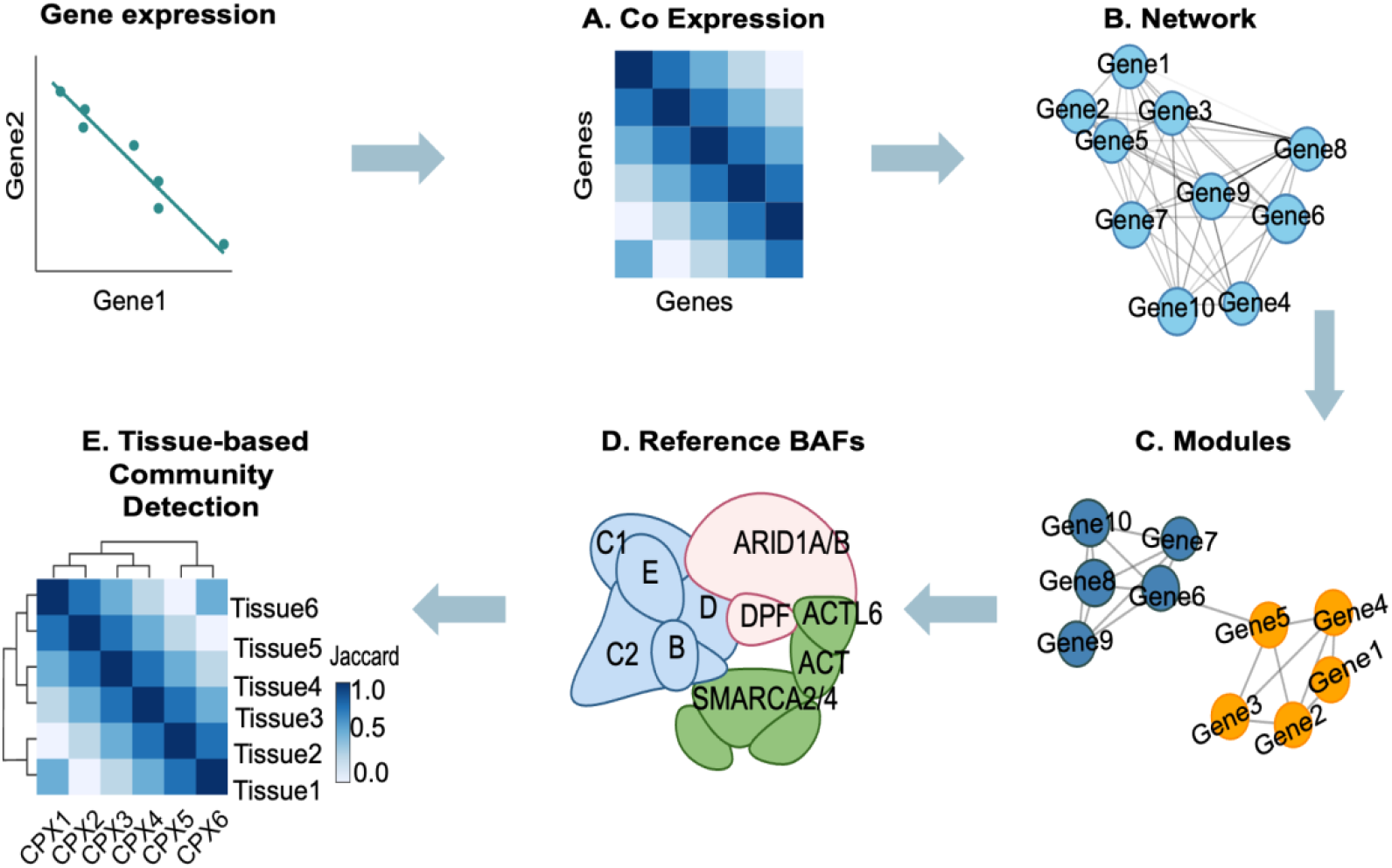
Overview of BAF gene co-expression network analysis.(A) Pairwise Pearson correlations among 30 core BAF genes were used as the measure of co-expression.(B) Construction of the co-expression network: correlations were rescaled to [0.1] and raised to a tissue-specific power *β* to achieve a scale-free topology fit (R^2^ ≥ 0.70).(C) Louvain module detection (*γ* = 1) was run 100× per tissue: only modules with ≥ 95 % consensus membership were retained.(D) Curated reference BAF subcomplex gene sets, annotated by tissue type (panel created in BioRender).(E) Jaccard similarity between tissue modules and reference complexes, with empirical p-vahies derived from 1 000 permutations, visualized as a clustered heatmap across tissues. BAF complexes, and endocrine categories.

### Co-Expression Network Construction

For each of the 49 GTEx tissue types, we calculated all pairwise Pearson correlation coefficients (r) between the 30 BAF-complex genes after applying a log_2_ (TPM + 1) transformation. Correlations whose absolute value exceeded √0.70 (|r| ≥ 0.837, corresponding to r^2^ ≥ 0.70) were set to zero, effectively down-weighing excessively strong, potentially redundant edges. Although one could apply a Benjamini–Hochberg correction across the 435 gene-pair tests per tissue—corresponding to significance thresholds of roughly |r| ≈ 0.30 in large-sample tissues and |r| ≈ 0.25 in smaller ones—such low cutoffs would produce overly dense networks in well-powered tissues and nearly empty ones in underpowered tissues. Because our goal was to recover a biologically meaningful, scale-free topology rather than simply flag statistically significant edges, we adopted the R^2^ ceiling approach. This preserves overall co-expression structure, prevents domination by a few ultra-strong correlations, and ensures consistency across tissues with varying sample sizes.

To maintain informative edge weights while promoting a scale-free network topology, we utilized WGCNA’s signed soft-thresholding method rather than strict binarization. For each tissue, we set a specific random seed before running WGCNA’s function to pick a soft threshold, testing a range of β values from 1 to 40. We selected the lowest β value achieving a scale-free topology fit of R^2^ ≥ 0.70; if no exponent satisfied this criterion, β was set to 6 by default.

Adjacency weights (*a*_*ij*_) were then calculated as:

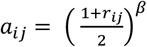

Here, *r*_*ij*_ represents the Pearson correlation between genes i and j (ranging from –1 to +1 after log_2_ transformation), and β is the chosen soft-thresholding exponent.

Adjacency values lower than 0.10 were discarded and self-loops were removed, and the largest connected component of at least five genes—the minimum number corresponding to known BAF sub-complexes—was retained for further analysis. We further enforced a minimum module size of five genes to define a community, for two reasons. First, our goal is to quantify coordinated changes to BAF composition and regulation. Second, because that is the smallest curated protein complex from our dataset.

### Module Detection

Community detection in each tissue-specific weighted graph was performed using the Louvain algorithm at the default resolution (γ = 1), optimizing weighted Newman–Girvan modularity (Q):

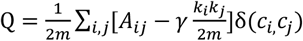

In this formula, *A*_*ij*_is the absolute adjacency between genes i and j; *k*_*i*_ is the weighted degree of gene i; m represents half of the total edge weight in the network; γ is the resolution parameter controlling community size; and *δ*(*c*_*i*,_ *c*_*j*_) equals 1 when genes i and j are in the same module, otherwise 0.

To enhance reproducibility, the Louvain algorithm was run 100 times per tissue, each with a unique random seed. A gene was assigned to a consensus community only if it appeared consistently (≥95% of the runs). Genes without stable community assignments were considered not clustered and module quality was evaluated by global modularity (Q) in each tissue.

Tissue-specific networks were visualized using fixed layouts in which node colored by consensus community membership and. Tissue-specific co-expression networks were visualized using a fixed node layout, ensuring that each gene occupies the same position across all tissues. This allows for direct comparison of network structure between tissues. Node colors indicate consensus community membership as determined by Louvain clustering.

This methodology generates robust and clearly defined gene modules, capturing nuanced co-expression patterns and facilitating subsequent functional and biological interpretations.

### Community Validation and Visualization

We evaluated the concordance between identified gene communities and 46 curated reference BAF protein complexes (**Figure 3**, Additional **Table S1**) across 45 tissues using the Jaccard Index as a metric. The Jaccard Index quantified shared gene memberships as: 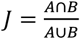, where A represents genes in the reference complexes and B represents genes in the detected community. Permutation testing (n = 1000) generated empirical p-values for the Jaccard Index by randomly selecting gene sets of equivalent size, assessing how frequently these permutations yielded higher or equal similarity than observed. A clustered heatmap of Jaccard indices between tissue-specific Louvain communities and reference BAF complexes was generated, annotated by endocrine versus non-endocrine tissue categories. Analyses were performed in RStudio Server on MCW’s RCC using R v4.4.2 and the packages WGCNA and igraph.

**Figure 3.**
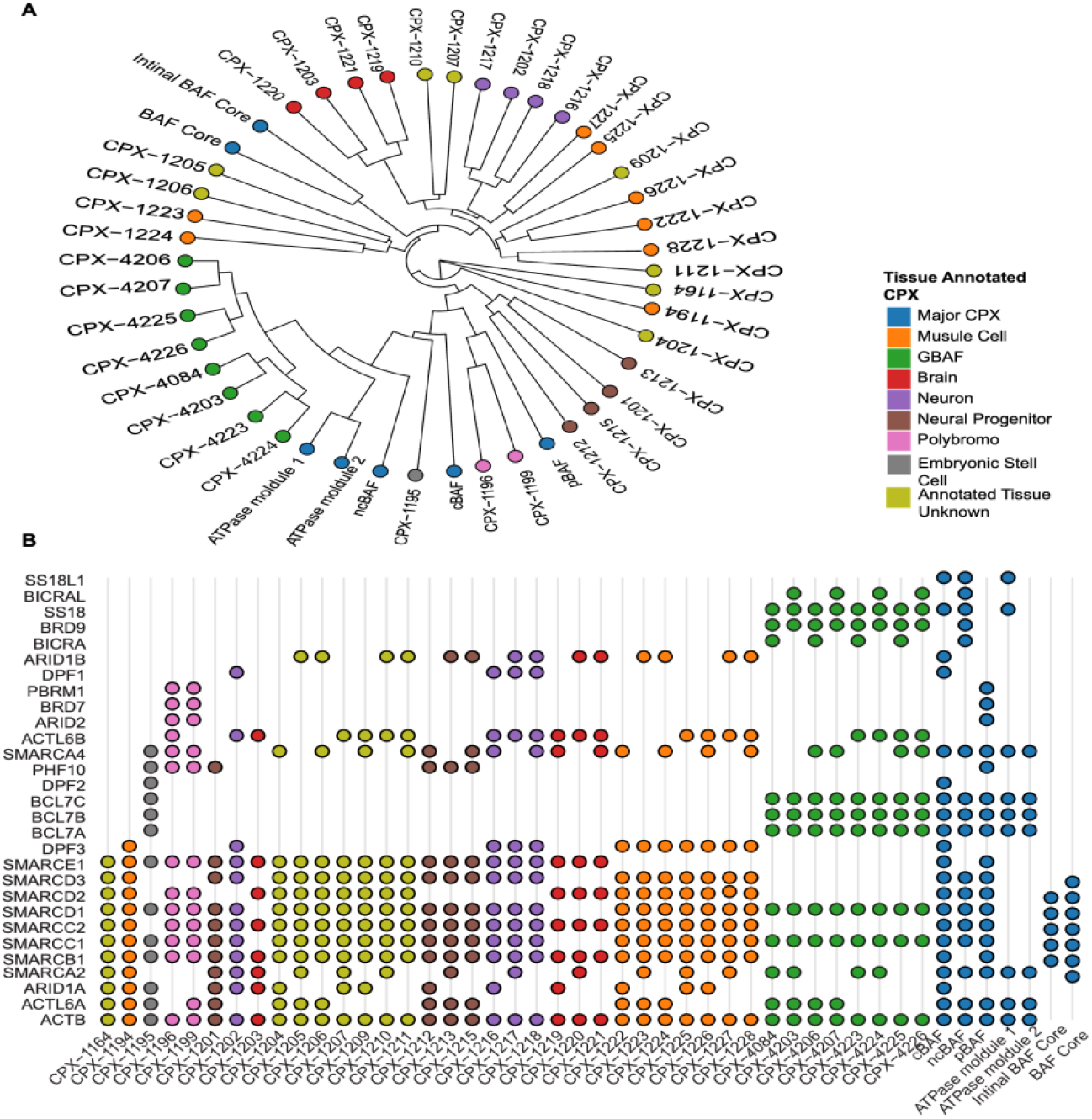
Diverse BAF complexes are indexed in high-throughput databases, indicating tissue-specific configurations. A) A phylogenetic dendrogram shows that most body tissues align with a reference BAF composition, yet many display distinct patterns from the three most well-defined versions. B) A tanghulu-style dot matrix (a matrix where colored dots represent the presence or absence of BAF complex genes, visually resembling candied fruit on a stick) displays the reference BAF complex gene composition by tissue type, highlighting shared core module components and a diverse array of variable components. Of the 46 reference BAF protein complexes, the major complexes—including cBAF, ncBAF, pBAF, BAF cores, initial BAF cores, ATPase module 1, and ATPase module 2—were sourced from Mashtalir et al. [2], while the remaining 39 reference complexes were obtained from the Complex Portal database [25].

## Results

### Descriptive Analysis of GTEx data

We classified transcriptomic samples into four endocrine categories—supporting, secondary endocrine, primary endocrine, and others—to assess sample representation across endocrine tissue groups (**Figure 4**). The endocrine “other” category supplies 11,663 of 19,452 samples (59.96%); the endocrine primary adds 4,254 samples (21.87%); the endocrine secondary tissues contribute 2,285 samples (11.75%); and endocrine supporting tissues were the fewest with 1,250 samples (6.43%). Sample sizes per tissue ranged from 181 (brain, amygdala) to 818 (muscle, skeletal), with a median of 277 samples. This unbalanced sample distribution underscores the importance of downstream methods, such as soft thresholding and consensus community detection, that accommodate variable sampling depth across tissue types.

**Figure 4.**
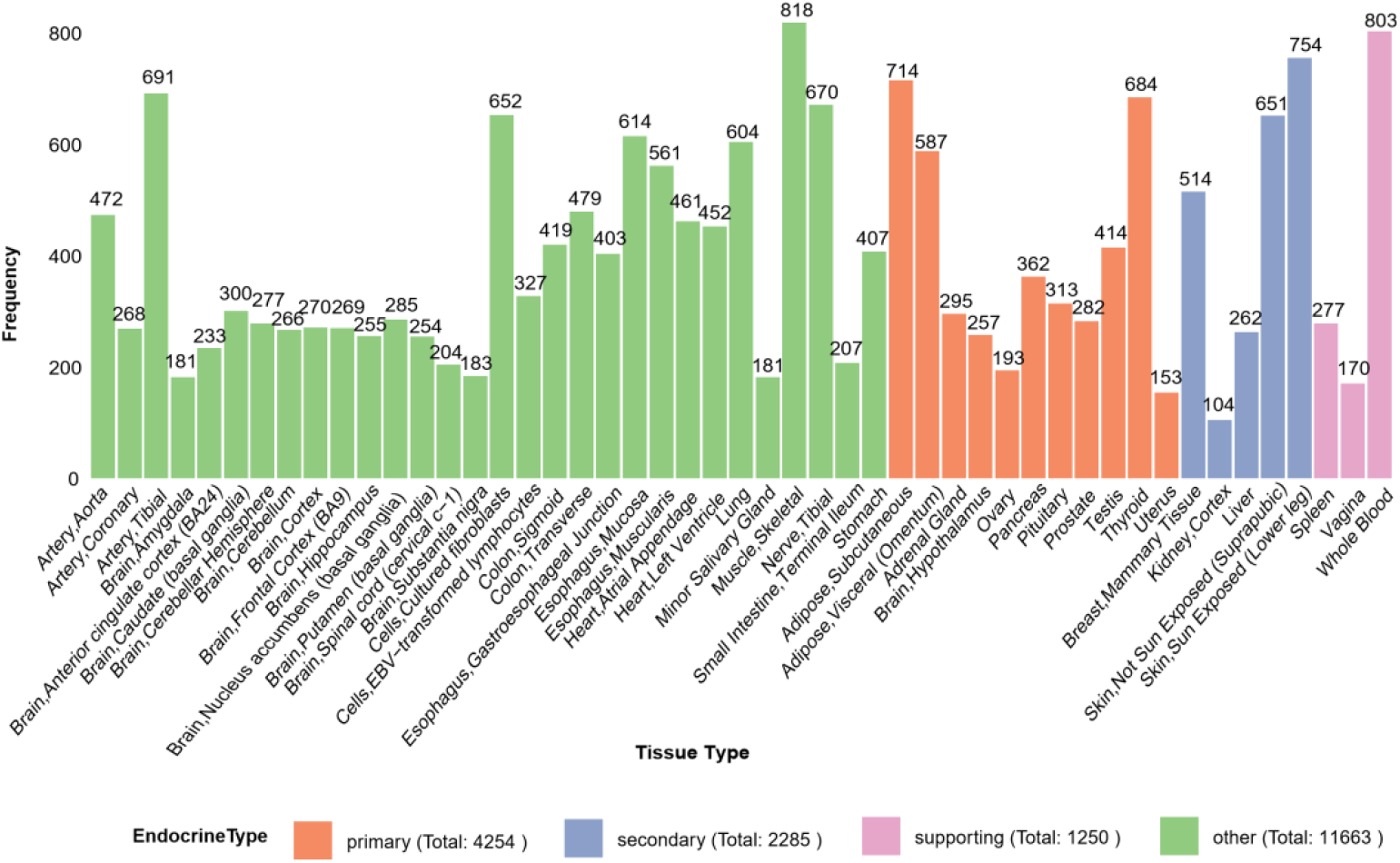
Sample distribution across GTEx tissues by endocrine annotation. Bar plot showing the distribution of RNA-seq samples across tissues after filtering for ≥100 samples per tissue. Tissues are grouped and color-coded by endocrine category: primary endocrine (green), secondary endocrine (orange), supporting (blue), and other (purple). Tissue names and sample counts are indicated along the axes. Category definitions are provided in the Methods

### BAF genes co-expression network analysis

Of the 49 tissues that passed the initial sample-size filter, four—Vagina, Lung, Spleen, and Colon, Sigmoid— yielded no modules and were excluded from further analysis. In Lung and Spleen, no edges remained after soft-thresholding and pruning, resulting in empty networks. In Vagina and Colon, Sigmoid, only small, disconnected network containing fewer than five genes remained, failing to meet the requirement that a module include at least five genes. Consequently, co-expression networks were built for 45 tissues. The fitted soft-threshold exponent β ranged from 1 to 35 (median = 6, mean = 8.3). Global modularity (Q) values spanned –0.019 to 0.41 (median = 0.047, mean = 0.080), indicating modest but positive community structure on average. On balance, these β and Q distributions reveal generally modest yet biologically meaningful community structure across tissues.

### Community Validation and Visualization

**Tables 1.1–1.5** present only those tissue–complex pairs from Additional **Table S2** that met the criteria of Jaccard Index > 0.4 or P-Value < 0.05. A full comparison across all 45 tissue types and 46 reference BAF complexes is available in Additional **Table S2** in the Supplementary Document.

**Table 1.1:**
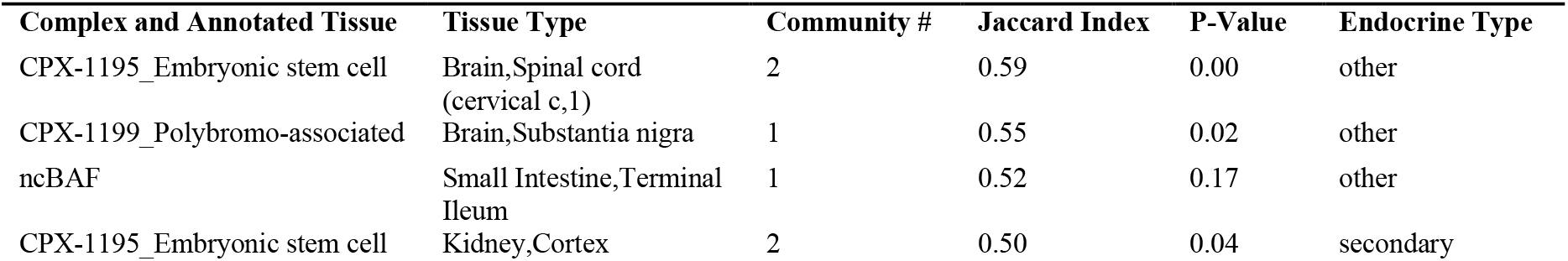
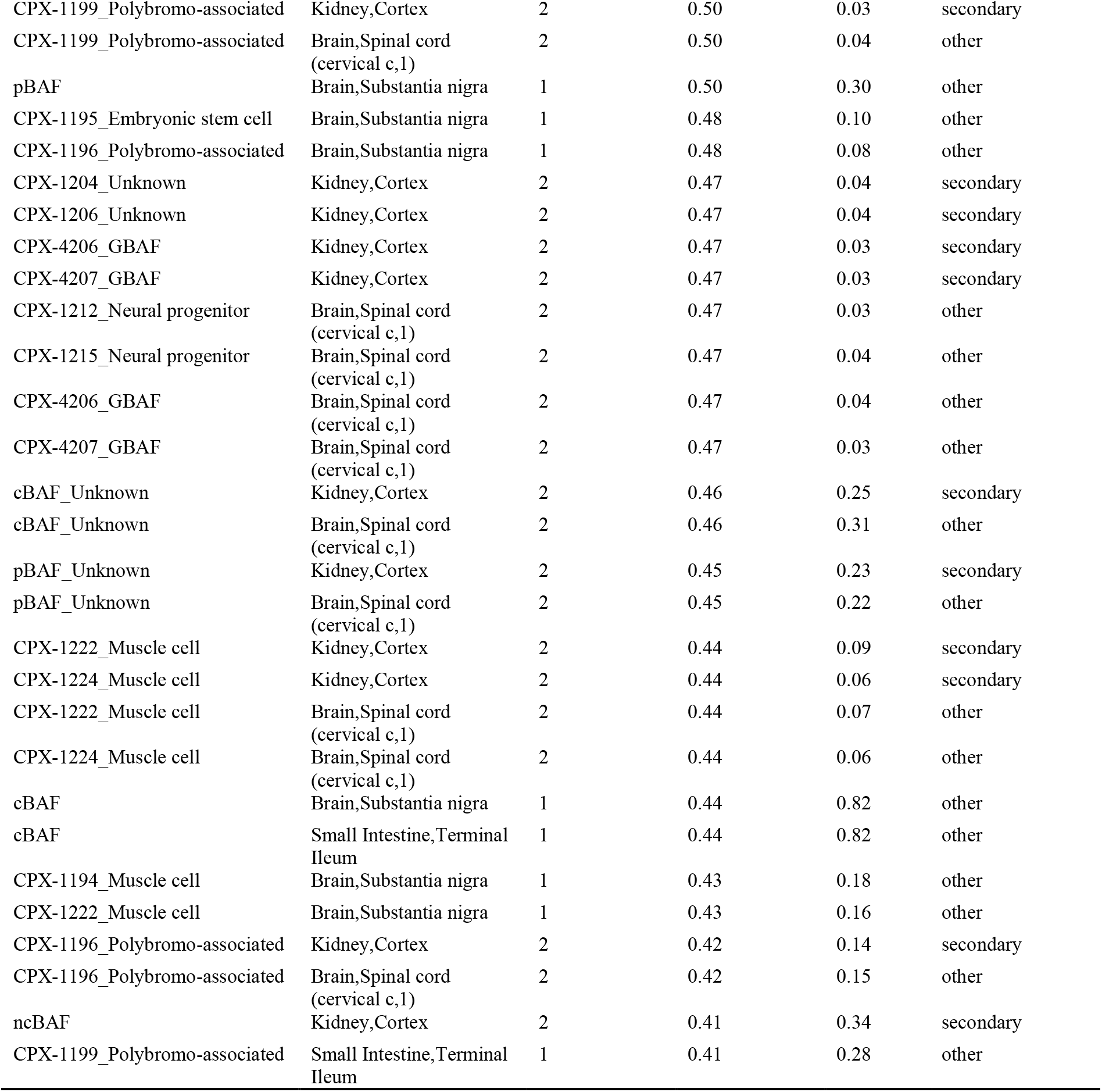
Block 1a Cluster. Captures Embryonic, Neural, and Secondary Kidney Tissues

The hierarchical clustering visualization between tissue-specific Louvain communities and reference BAF subcomplex gene sets—annotated by endocrine category—included only tissues with moderate-to-high overlap (Jaccard Index ≥ 0.3) to ensure clarity and streamlined interpretation.

Hierarchical clustering of the resulting similarity matrix **(Figure. 5)** revealed four discrete tissue clusters, termed Blocks 1a,1b,1c,2 and 3. Each Block is distinguished by a characteristic pattern of complex overlap and endocrine classification.

**Figure 5.**
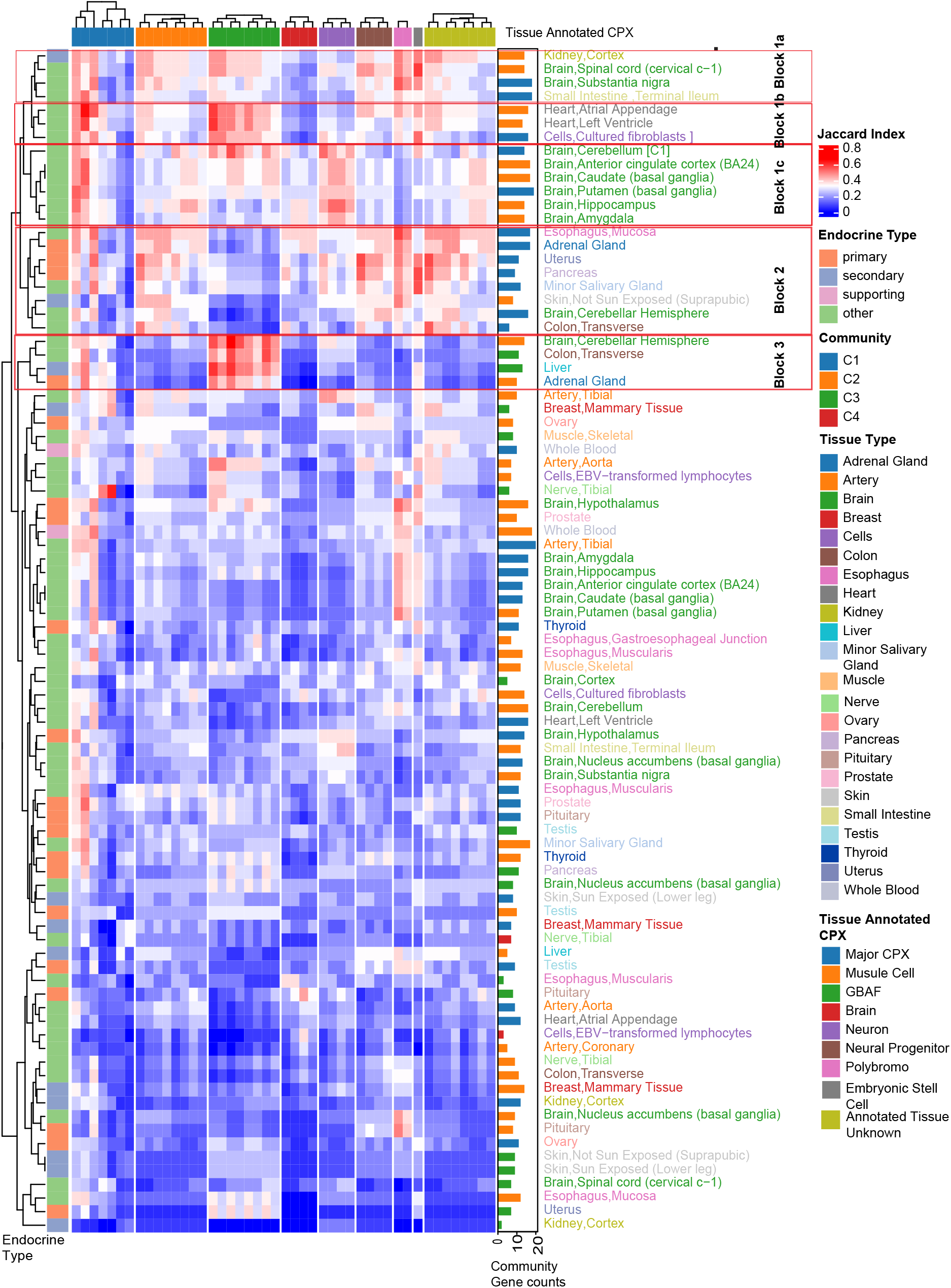
Heatmap of Jaccard similarity between GTEx tissue-specific BAF communities (rows) and curated CPX reference subcomplexes (columns). Columns are grouped and ordered by subcomplex category based on tissue annotation: **Major CPX:** cBAF, ncBAF, pBAF, ATPase module_1, ATPase module_2, BAF Core, Initial BAF Core. **Muscle cell:** CPX-1222, CPX-1224, CPX-1194, CPX-1223, CPX-1226, CPX-1228, CPX-1227, CPX-1225. **GBAF:** CPX-4206, CPX-4207, CPX-4225, CPX-4226, CPX-4084, CPX-4203, CPX-4223, CPX-4224. **Brain:** CPX-1219, CPX-1221, CPX-1220, CPX-1203. **Neuron:** CPX-1216, CPX-1218, CPX-1217, CPX-1202. **Neuronal progenitor:** CPX-1212, CPX-1215, CPX-1201, CPX-1213. **Polybromo-associated:** CPX-1199, CPX-1196. **Embryonic:** CPX-1195. **Unknown tissue annotation:** CPX-1204, CPX-1206, CPX-1164, CPX-1205, CPX-1209, CPX-1211, CPX-1210, CPX-1207. Rows represent the 49 GTEx tissues, with side color bars indicating endocrine classification (primary, secondary, supporting, other) and adjacent bar graphs showing community gene counts. Cell color encodes the Jaccard index (degree of gene overlap) between each tissue’s Louvain community and the reference gene set. The similarity matrix revealed four discrete tissue clusters, termed Blocks 1a, 1b, 1c, 2, and 3. Additional **Figure S5** (supplementary to **Figure 5**) presents the same data structure and layout, but includes a column for the CPX ID associated with each reference BAF complex.

### Function-Oriented Module Analysis Reveals Endocrine and Non-Endocrine patterning Block 1a Captures Embryonic, Neural, and Secondary Kidney Tissues

Block 1a comprises embryonic stem cell–associated (CPX-1195), cBAF, polybromo-associated (CPX-1199), and neural progenitor chromatin-remodeling modules, with prominent overlap in the spinal cord cervical segment, substantia nigra (endocrine “other”), and kidney cortex (endocrine “secondary”). Notably, the embryonic stem cell module CPX-1195 shows high similarity with the spinal cord (Jaccard index = 0.59, P-value < 0.001), as shown in **Figure 5, Figure 6A, and Table 1.1 (Block 1a Cluster).** These underscores shared embryonic/neural-like chromatin mechanisms that extend into secondary renal tissues.

**Figure 6.**
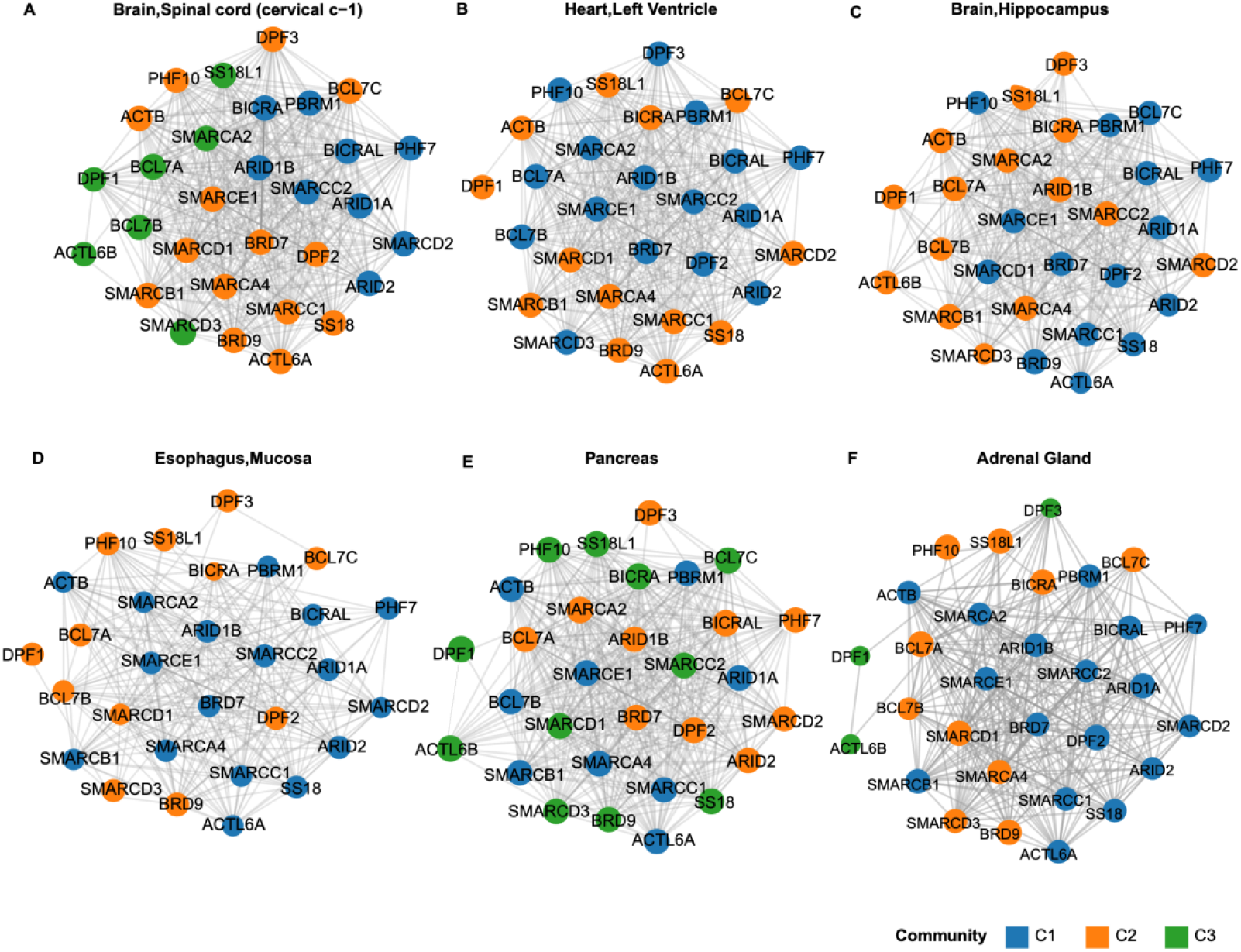
Representative Louvain consensus community networks reveal differences in BAF gene regulation across tissues. Tissue specific co-expression networks of 30 BAF complex genes are shown for one or two representative tissues from each major cluster (Blocks 1a, 1b, 1c, 2, and 3, as defined in **Figure 5**). Networks are displayed using a fixed arrangement so that each gene occupies the same position across all panels. Edges denote adjacency weights > 0.10 (after signed soft thresholding), and nodes are colored by their Louvain consensus community membership. This positioning highlights how each tissue assembles distinct BAF subcomplex modules.

### Block 1b Highlights Cardiac, Cerebellar, and Supporting Fibroblast Tissues with GBAF Modules

Block 1b features cardiac tissues (atrial appendage, left ventricle; endocrine “other”), cerebellum (endocrine”other”), and cultured fibroblasts (endocrine “supporting”). These tissues exhibit significant enrichment for non-canonical BAF (ncBAF) and GBAF modules, particularly in the cardiac atrial appendage (ncBAF: Jaccard = 0.60, P-value = 0.01), and left ventricle (GBAF CPX-4206: Jaccard = 0.60, P-value < 0.001) which shown in **Figure 5, Figure 6B, and Table 1.2 - Block 1b Cluster.** The cerebellum shows robust alignment with GBAF (CPX-4225: Jaccard = 0.56, P = 0.01) and neuron-specific modules (CPX-1216: Jaccard = 0.53, P = 0.01), whereas fibroblasts align with embryonic and GBAF modules. These patterns reflect conserved chromatin remodeling mechanisms essential to cardiac, neural, and stromal tissue identities.

**Table 1.2:** Block 1b Cluster. Highlights Cardiac, Cerebellar, and Supporting Fibroblast Tissues with GBAF Modules

| Complex and Annotated Tissue | Tissue Type | Community # | Jaccard Index | P-Value | Endocrine Type |
| --- | --- | --- | --- | --- | --- |
| ncBAF | Heart,Atrial Appendage | 2 | 0.65 | 0.01 | other |
| CPX-4206_GBAF | Heart,Atrial Appendage | 2 | 0.59 | 0.00 | other |
| pBAF | Heart,Atrial Appendage | 2 | 0.55 | 0.07 | other |
| CPX-4084_GBAF | Heart,Atrial Appendage | 2 | 0.50 | 0.02 | other |
| CPX-4207_GBAF | Heart,Atrial Appendage | 2 | 0.50 | 0.03 | other |
| CPX-4225_GBAF | Heart,Atrial Appendage | 2 | 0.50 | 0.02 | other |
| cBAF | Heart,Atrial Appendage | 2 | 0.48 | 0.41 | other |
| CPX-1195_Embryonic stem cell | Heart,Atrial Appendage | 2 | 0.45 | 0.12 | other |
| CPX-1199_Polybromo-associated | Heart,Atrial Appendage | 2 | 0.45 | 0.14 | other |
| CPX-1204_Unknown | Heart,Atrial Appendage | 2 | 0.42 | 0.12 | other |
| CPX-1206_Unknown | Heart,Atrial Appendage | 2 | 0.42 | 0.10 | other |
| CPX-1212_Neural progenitor | Heart,Atrial Appendage | 2 | 0.42 | 0.12 | other |
| CPX-1215_Neural progenitor | Heart,Atrial Appendage | 2 | 0.42 | 0.12 | other |
| CPX-4203_GBAF | Heart,Atrial Appendage | 2 | 0.42 | 0.12 | other |
| CPX-4223_GBAF | Heart,Atrial Appendage | 2 | 0.42 | 0.10 | other |
| CPX-4226_GBAF | Heart,Atrial Appendage | 2 | 0.42 | 0.09 | other |
| CPX-1222_Muscle cell | Heart,Atrial Appendage | 2 | 0.40 | 0.20 | other |
| CPX-1224_Muscle cell | Heart,Atrial Appendage | 2 | 0.40 | 0.22 | other |
| CPX-4206_GBAF | Heart,Left Ventricle | 2 | 0.60 | 0.00 | other |
| ncBAF | Heart,Left Ventricle | 2 | 0.58 | 0.01 | other |
| CPX-4084_GBAF | Heart,Left Ventricle | 2 | 0.50 | 0.02 | other |
| CPX-4207_GBAF | Heart,Left Ventricle | 2 | 0.50 | 0.01 | other |
| CPX-4225_GBAF | Heart,Left Ventricle | 2 | 0.50 | 0.02 | other |
| cBAF | Heart,Left Ventricle | 2 | 0.48 | 0.14 | other |
| CPX-4225_GBAF | Brain,Cerebellum | 1 | 0.56 | 0.01 | other |
| CPX-1216_Neuron | Brain,Cerebellum | 1 | 0.53 | 0.01 | other |
| CPX-4206_GBAF | Brain,Cerebellum | 1 | 0.47 | 0.02 | other |
| CPX-4223_GBAF | Brain,Cerebellum | 1 | 0.47 | 0.03 | other |
| CPX-4226_GBAF | Brain,Cerebellum | 1 | 0.47 | 0.04 | other |
| cBAF | Brain,Cerebellum | 1 | 0.46 | 0.32 | other |
| CPX-1202_Neuron | Brain,Cerebellum | 1 | 0.44 | 0.09 | other |
| CPX-1218_Neuron | Brain,Cerebellum | 1 | 0.44 | 0.07 | other |
| CPX-1226_Muscle cell | Brain,Cerebellum | 1 | 0.44 | 0.08 | other |
| CPX-1195_Embryonic stem cell | Brain,Cerebellum | 1 | 0.42 | 0.14 | other |
| ncBAF | Brain,Cerebellum | 1 | 0.41 | 0.34 | other |
| CPX-1195_Embryonic stem cell | Cells,Cultured fibroblasts | 1 | 0.53 | 0.03 | other |
| CPX-4206_GBAF | Cells,Cultured fibroblasts | 1 | 0.50 | 0.02 | other |
| CPX-4207_GBAF | Cells,Cultured fibroblasts | 1 | 0.50 | 0.02 | other |
| cBAF | Cells,Cultured fibroblasts | 1 | 0.48 | 0.42 | other |
| pBAF | Cells,Cultured fibroblasts | 1 | 0.48 | 0.26 | other |
| CPX-4084_GBAF | Cells,Cultured fibroblasts | 1 | 0.42 | 0.12 | other |
| CPX-4203_GBAF | Cells,Cultured fibroblasts | 1 | 0.42 | 0.10 | other |
| CPX-4225_GBAF | Cells,Cultured fibroblasts | 1 | 0.42 | 0.11 | other |
| CPX-4226_GBAF | Cells,Cultured fibroblasts | 1 | 0.42 | 0.11 | other |
| CPX-1216_Neuron | Cells,Cultured fibroblasts | 1 | 0.40 | 0.21 | other |
| CPX-1222_Muscle cell | Cells,Cultured fibroblasts | 1 | 0.40 | 0.20 | other |

### Block 1c Defines Neuron-Specific Remodeling in Cortical and Subcortical Brain Regions

Block 1c encompasses the anterior cingulate cortex, caudate, putamen, hippocampus, and amygdala (“other”), and is primarily associated with neuron-specific BAF modules. For example, the hippocampus shows robust alignment with neuron-specific BAF module CPX-1218 (Jaccard index = 0.53, P-value = 0.01), as shown in **Figure 5, Figure 6C, and Table 1.3-Block 1c Cluster).** Additional associations are seen with ncBAF and cBAF modules. This pattern highlights specialized, neuron-centric chromatin remodeling mechanisms essential for cognitive and emotional functions in these brain regions.

**Table 1.3:**
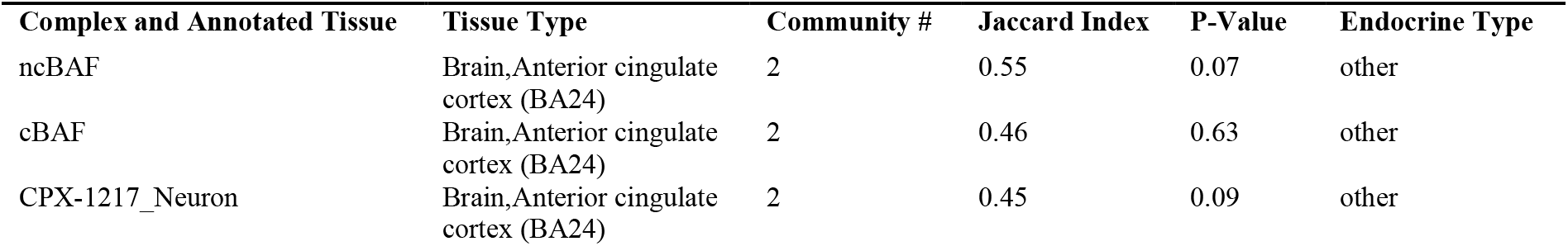
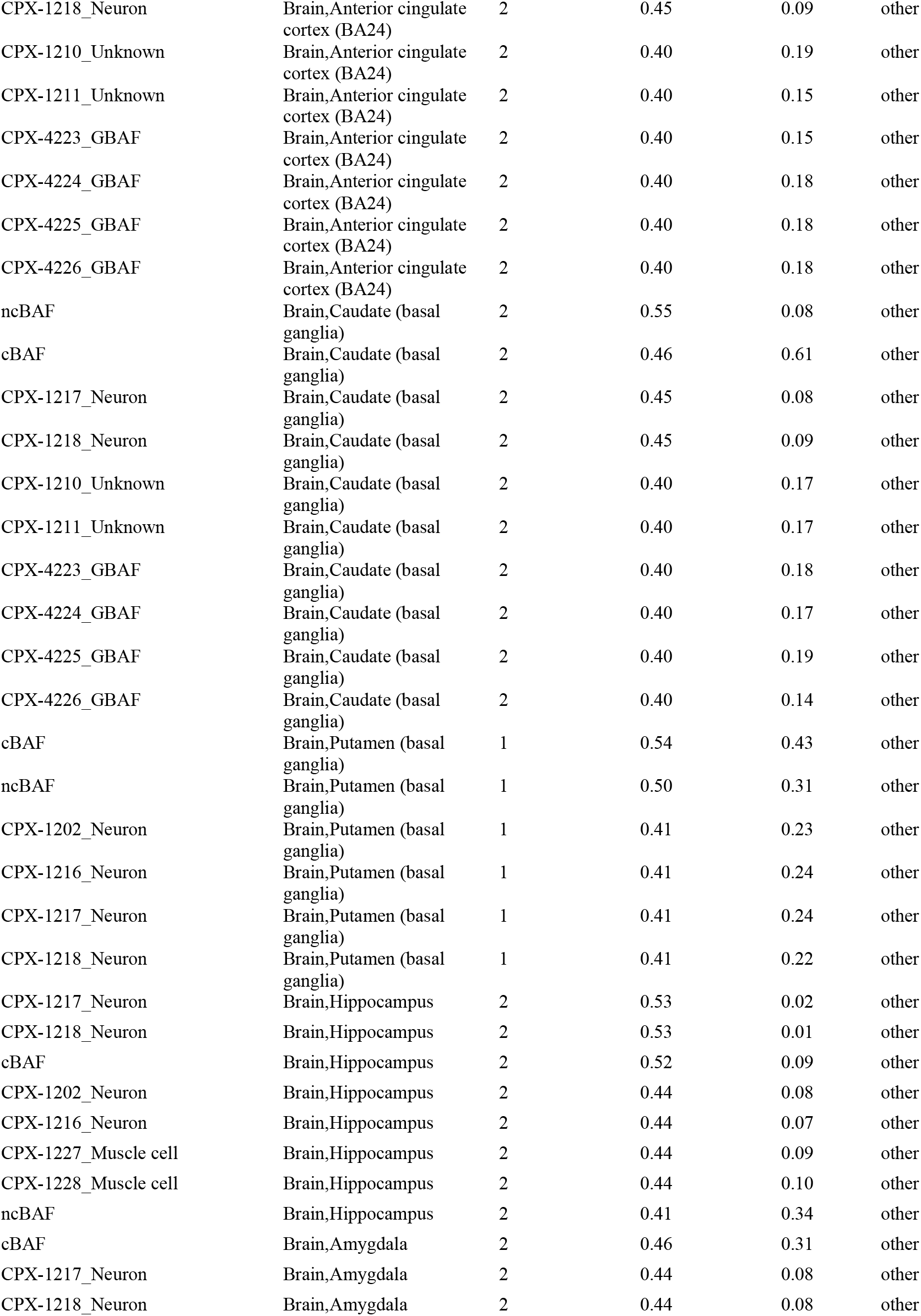
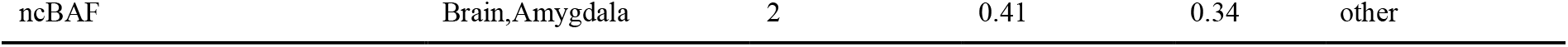
Block 1c Cluster. Defines Neuron-Specific Remodeling in Cortical and Subcortical Brain Regions

### Block 2 Represents Endocrine and Mucosal Chromatin Remodeling Patterns

Block 2 includes primary endocrine tissues (adrenal gland, pancreas, uterus) and mucosal epithelia (esophagus, colon, salivary gland, skin). These tissues are enriched for polybromo-associated and muscle-related modules, with strong interactions observed in esophageal mucosa (CPX-1199: Jaccard 0.58, P-value = 0.01 shown in **Figure 5, Figure 6D, and Table 1.2: Block 2 Cluster**), adrenal gland, and pancreas. This reflects key chromatin regulation supporting both endocrine secretion and mucosal barrier function. The pancreas (CPX-1195: Jaccard index 0.57, P-value < 0.001**; Figure 5, Figure 6E, Table 1.4-Block 2 Cluster**) and tissues such as the minor salivary gland and uterus also show shared enrichment in embryonic stem cell-related modules.

**Table 1.4:**
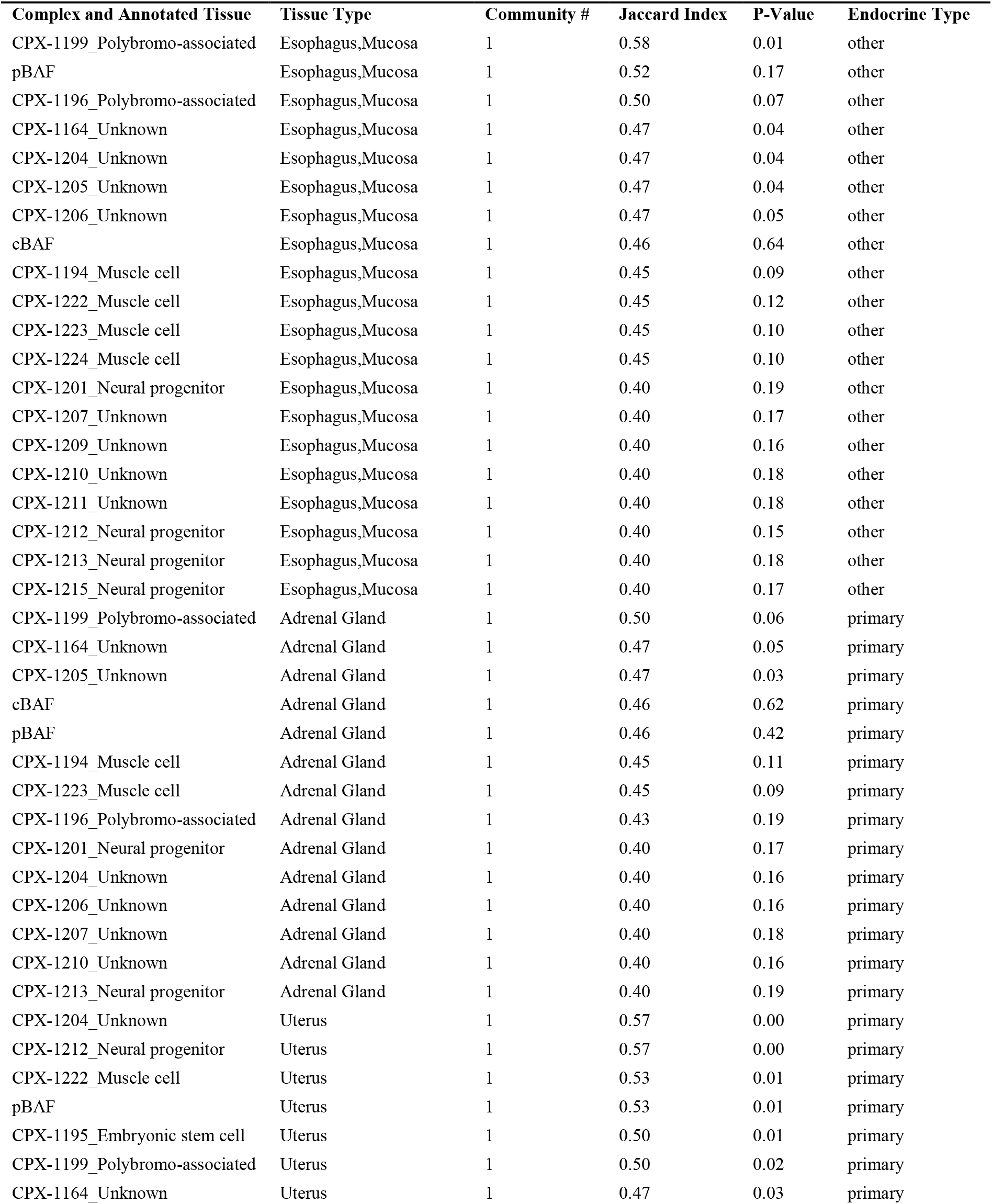
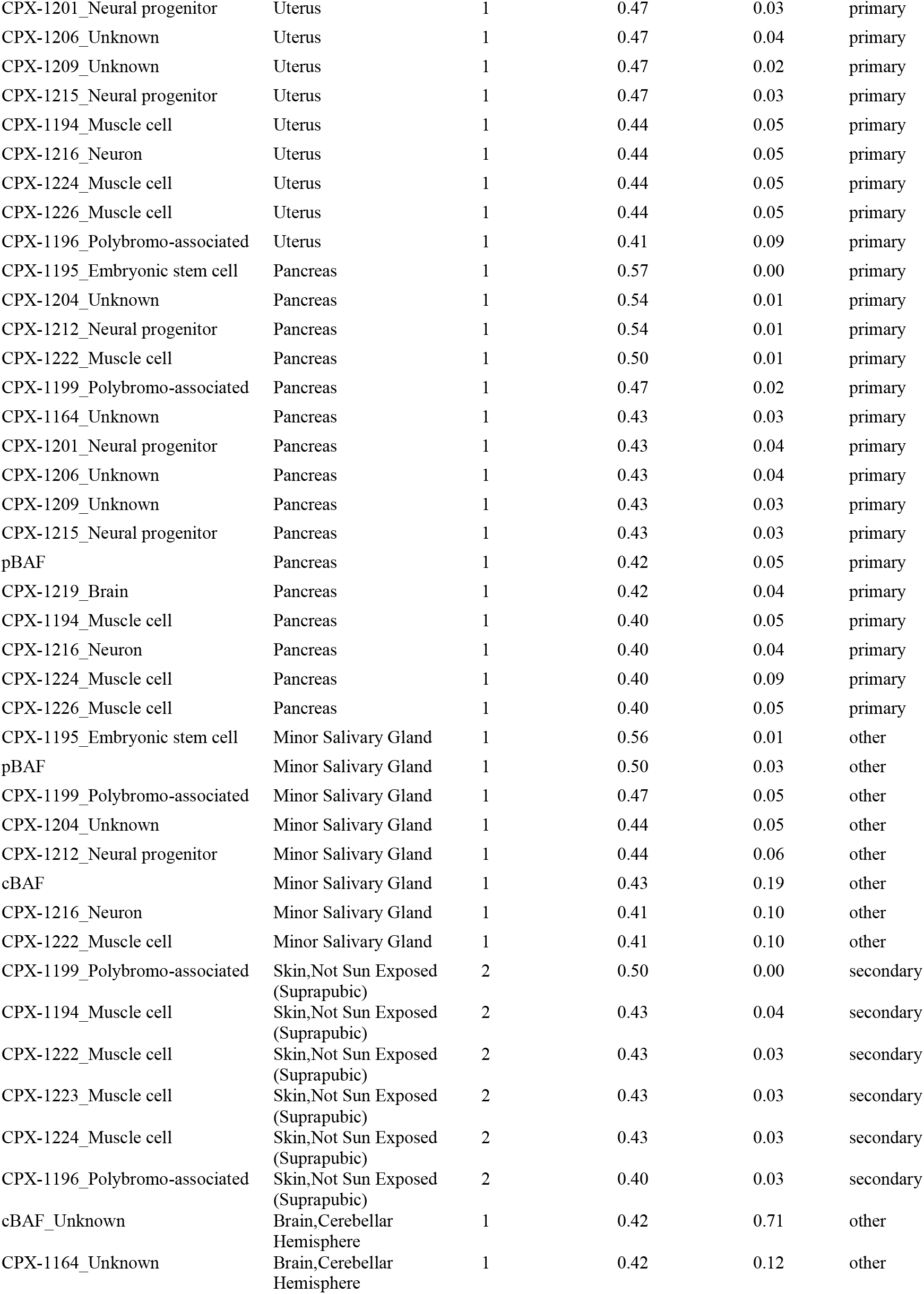
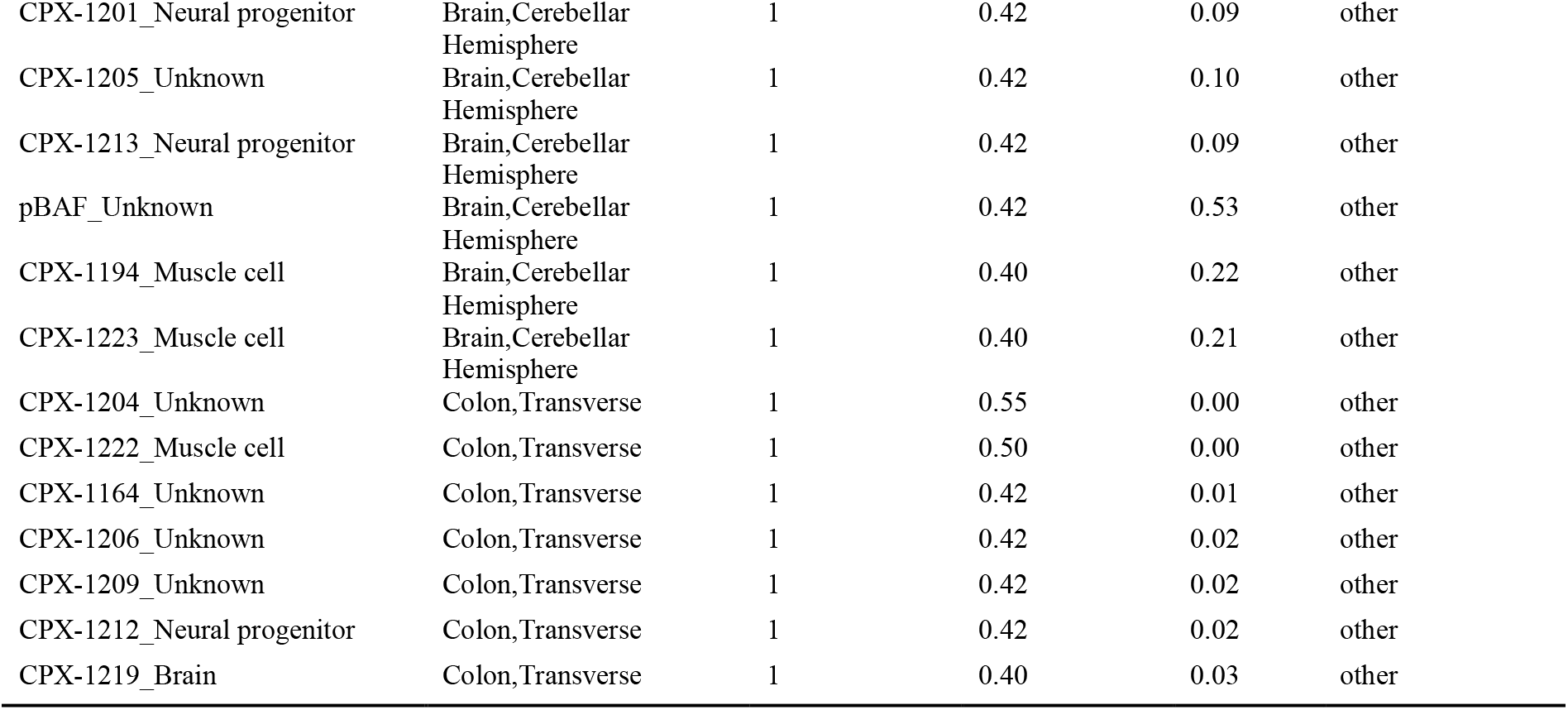
Block 2 Cluster. Represents Endocrine and Mucosal Chromatin Remodeling Patterns

### Block 3 Highlights GBAF Remodeling Across Cerebellar, Gastrointestinal, and Hepatic Tissues

Block 3 encompasses the cerebellar hemisphere, colon (transverse), liver, and adrenal gland, each showing strong associations with GBAF complexes. Notable examples include CPX-4225 in cerebellum (Jaccard 0.67, P < 0.001), CPX-4223 and CPX-4225 in colon (Jaccard 0.57, P < 0.001), as well as ncBAF in adrenal gland (Jaccard index 0.50, P = 0.02), as shown **in Figure 5, Figure 6F, and Table 1.5-Block 3 Cluster**. These findings indicate specialized, GBAF-driven chromatin regulation in metabolic, endocrine, and neuronal tissue contexts.

**Table 1.5:**
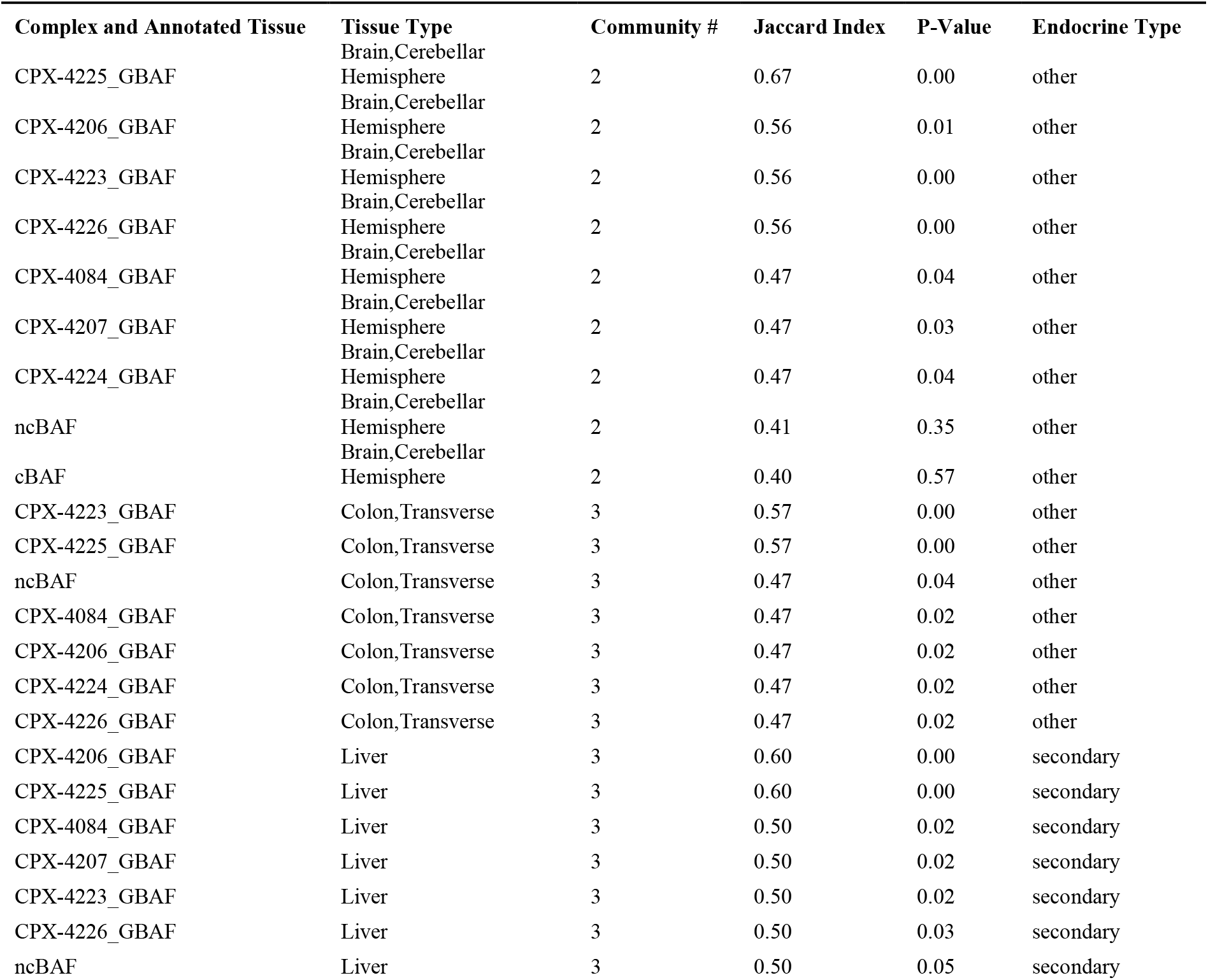
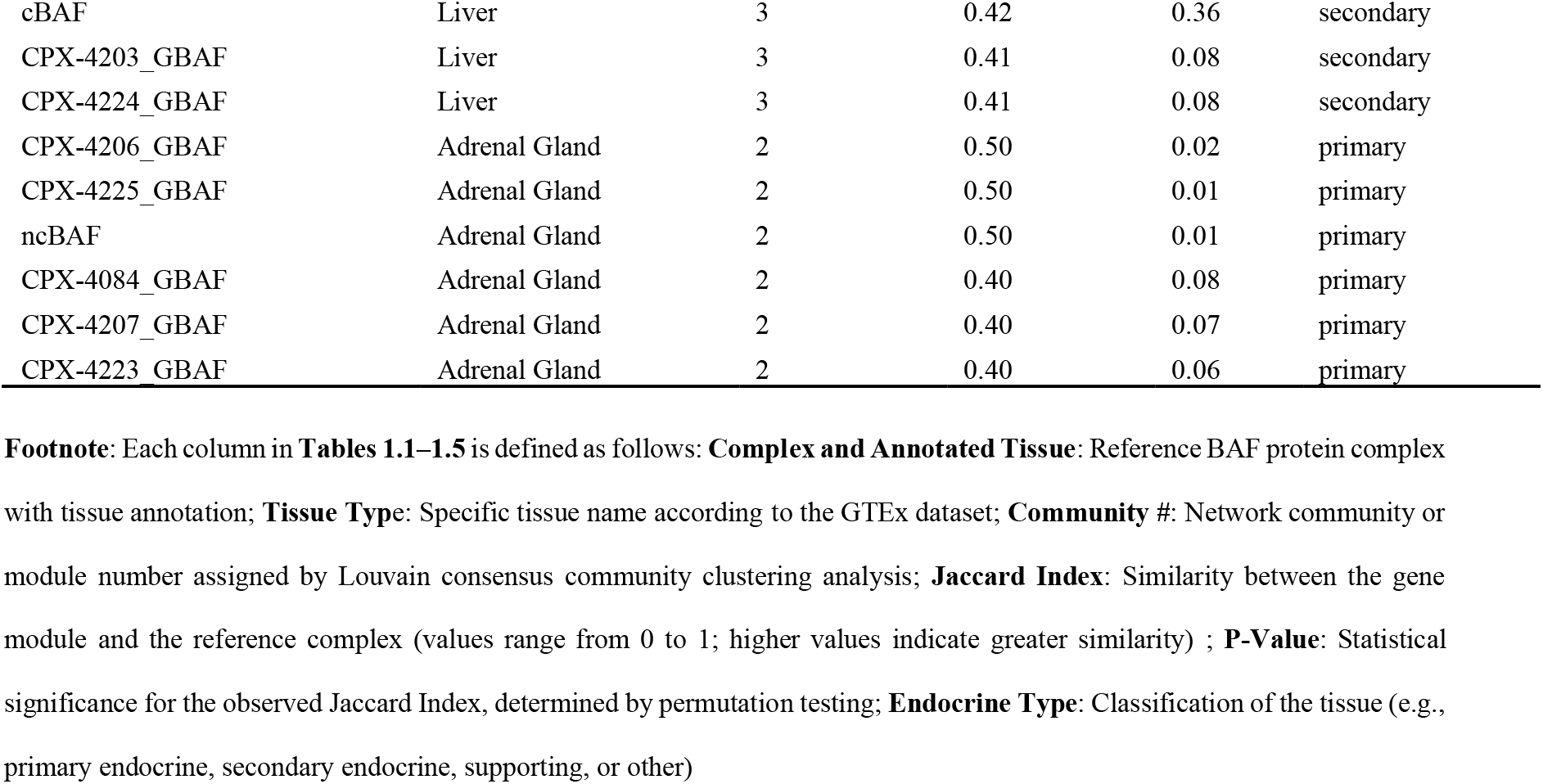
Block3 Cluster. Highlights GBAF Remodeling Across Cerebellar, Gastrointestinal, and Hepatic Tissues

## Discussion

We have taken first to our knowledge, data-driven large-scale analysis for the information about differential regulation of BAF complexes across human endocrine and non-endocrine tissues. Our analysis reveals that BAF complex subunits partition into gene co-expression communities whose reuse of chromatin remodeling modules is strikingly tissue-specific rather than tissue-agnostic or random. Structural and contractile tissues consistently indicate canonical cBAF/pBAF scaffolds (e.g., ARID1A/SMARCC1, PBRM1/ARID2), reflecting a “housekeeping” chromatin architecture essential for barrier integrity, extracellular-matrix organization and contractility. In the central nervous system, gene communities align almost exclusively with neuron-specific nBAF subunits (ACTL6B, SS18L1, DPF1) and neural progenitor modules, underscoring the precision of chromatin remodeling required for synaptic plasticity and mature neuronal identity. Endocrine and gastrointestinal epithelia may rely on what has been described as a smooth muscle–like BAF, to integrate motility and secretory function. Certainly, challenges of tissue dissection could relate to some of these patterns. We aim to deconvolute signals in the future, by repeating similar analysis in single-cell experiments. Finally, immune, stromal and barrier tissues preferentially engage non-canonical GBAF complexes (BRD9/BICRA), illustrating context-specific remodeling strategies.

A few important caveats qualify these conclusions. First, unsupervised methods such as WGCNA soft-thresholding, Louvain clustering, and hierarchical linkage, involve parameter choices (e.g., soft-threshold power, community resolution, and clustering metrics), that inevitably introduce differences between implementations.

To ameliorate this potential challenge, we performed sensitivity analyses across reasonable parameter ranges and confirmed that the major blocks (canonical BAF in structural tissues, nBAF in brain, pBAF in epithelia, GBAF in immune/barrier) in our results are robust to a range of parameter values. Alternative algorithms or thresholds could yield different partitions. Second, two genes, ACTL6B and DPF1, are rarely and lowly expressed in most tissues, leading to zero-inflation in their community profiles and limiting our ability to gauge their tissue-specific roles. Targeted proteomic or single-cell assays will be necessary to improve characterization of genes and proteins with their patterning. Third, GTEx samples derive from multiple sequencing centers with varied library protocols. We applied stringent batch correction before network construction and confirmed that no results block corresponds exclusively to a single center, residual technical variation may still influence co-expression patterns. Future studies using uniformly processed cohorts will help eliminate this potential confounding factor.

Beyond these caveats, our findings extend prior reports of BAF complex plasticity in development and disease. The well-documented npBAF→nBAF switch during neuronal lineage commitment is recapitulated in adult brain tissues, confirming the persistence of neuron-specific remodeling programs and supporting our premise that differences in BAF regulation can be inferred from large-scale data analysis [26–28]. Likewise, the enrichment of cBAF/pBAF in mesenchymal and epithelial tissues echoes their broad expression of ARID1A/ARID2 and SMARCA subunits in maintaining homeostatic transcriptional programs [17,29]. The discovery of tissue communities whose data patterns show negligible overlap across the existing 46 curated BAF complex definitions (e.g., reproductive glands, certain barrier epithelia and deep-brain subregions), highlights gaps in understanding (i.e., “complex-dark”) and points to possible novel or tissue-restricted remodeling complexes yet to be characterized.

Several limitations remain. Our study focused on 30 core BAF complex genes and 46 predefined modules. Expansion to include additional subunits, cofactors or lineage-specific interactors in future analyses, may refine the observed patterns. Bulk tissue RNA-seq averages signals across heterogeneous cell populations. Single-cell or spatial transcriptomics could reveal sharper community boundaries or microenvironment-specific remodeling events [30]. Finally, while the Jaccard index offers a straightforward measure of gene-set overlap, integrating quantitative measures of complex abundance or activity (e.g., ChIP-seq, proteomics) would strengthen functional inferences about remodeling dynamics.

In summary, BAF complex communities are systematically tailored to tissue identity, with predictable reuse of canonical, neuron-specific, smooth muscle–like and non-canonical modules. This modular deployment aligns with organ-specific functional demands and highlights underexplored “complex-dark” compartments as fertile ground for the discovery of novel epigenetic regulatory processes and therapeutic targets.

## Conclusions

Our integrative analysis points toward a non-random, modular deployment of BAF chromatin remodeling machinery across different human tissues. Weighted co-expression communities of BAF complex subunits recapitulate known remodeling modules in a highly tissue-specific manner: canonical cBAF/pBAF show strong co-regulation signals in structural and contractile organs; neuron-specific nBAF programs define central nervous system modules; endocrine and smooth-muscle–like epithelia demonstrate BAF signatures where muscle-cell reference BAFs are tightly co-regulated; and immune, stromal, and barrier compartments preferentially modularize non-canonical GBAF complexes. By contrast, a subset of reproductive and barrier tissues, by contrast, remain “complex-dark,” pointing to novel or specialized configurations of remodelers yet to be characterized, or the unique co-regulatory programs active in these tissues. Our findings underscore that chromatin remodeling is systematically tailored to fulfill organ-specific functional demands and highlight underexplored tissues as promising venues for the discovery of new epigenetic regulators.

## Supporting information

Supplymentary files explaination

## Data availability

The dataset used during the current study is available from GTEx bulk RNA-seq data (V10 release) which is open data access that can be downloaded from the GTEx portal (https://gtexportal.org/home/). GTEx_Analysis_2022-06-06_v10_RNASeQCv2.4.2_gene_median_tpm.gct sequencing data GTEx_Analysis_v10_Annotations_SampleAttributesDS.txt includes detailed clinical characteristics of patients.

## Declaration of interest, Funding and Acknowledgements

### Author contribution statement

Conception and design: M.T. Zimmermann, X. Dong. Development of methodology: Xong, M.T. Zimmermann. Acquisition of data: X.Dong, M.T. Zimmermann. Analysis of data: X.Dong. Interpretation of data: X.Dong, N. Haque, J. Wagenknecht. Writing, review, and/or revision of the manuscript: X.Dong, M.T. Zimmermann.

## Acknowledgements

We thank Dr.s Colbie Reed and William Hogan at MCW for their assistance with data curation.

