## Supplementary figures and images for "Tissue-Specific Co-Expression Patterns of BAF Complexes Provide Regulatory Insights Across Human Tissues with Implications for Endocrine and Non-Endocrine Functions"

### Supplymemtal Figure S1

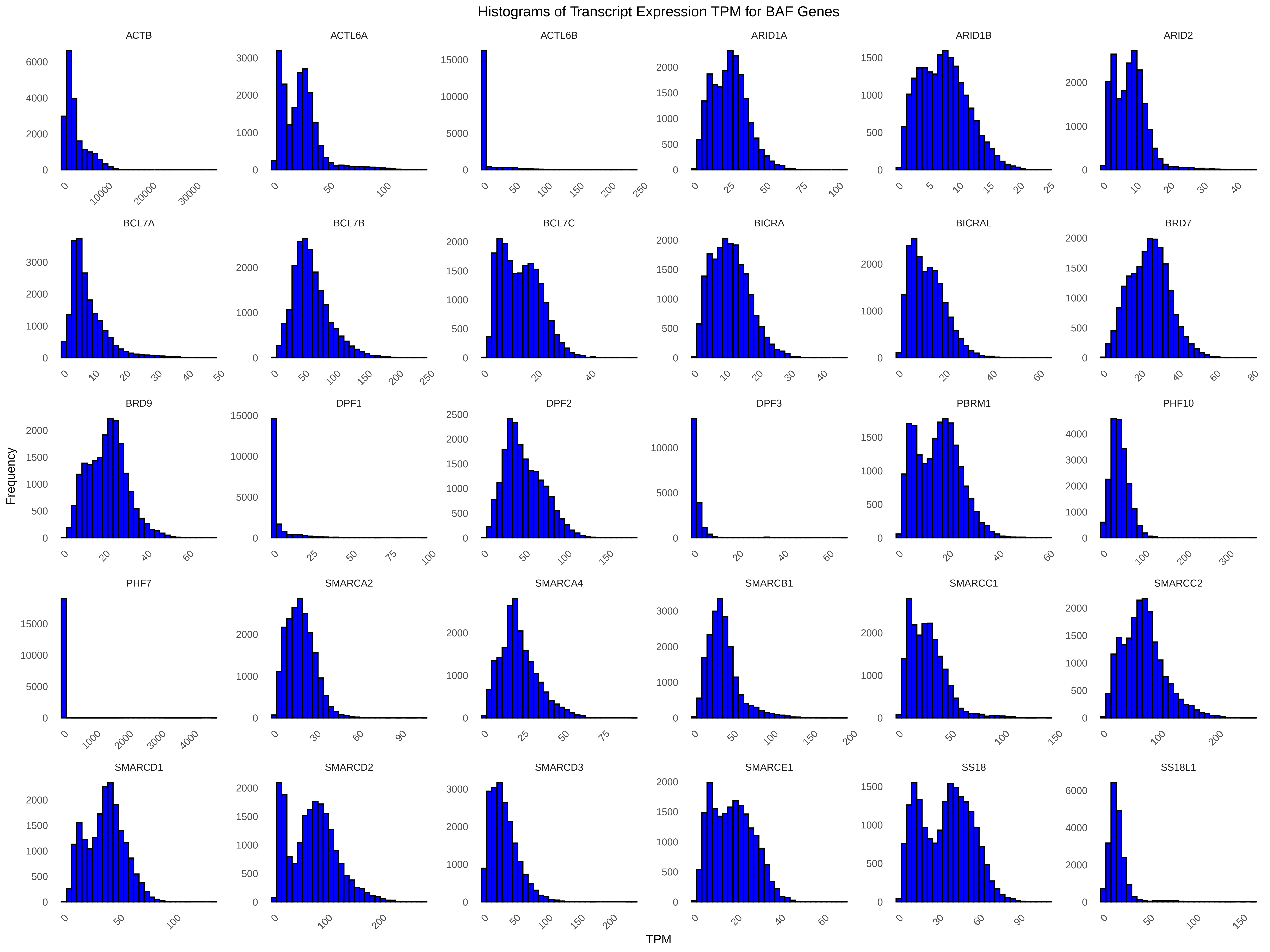

### Supplymemtal Figure S2

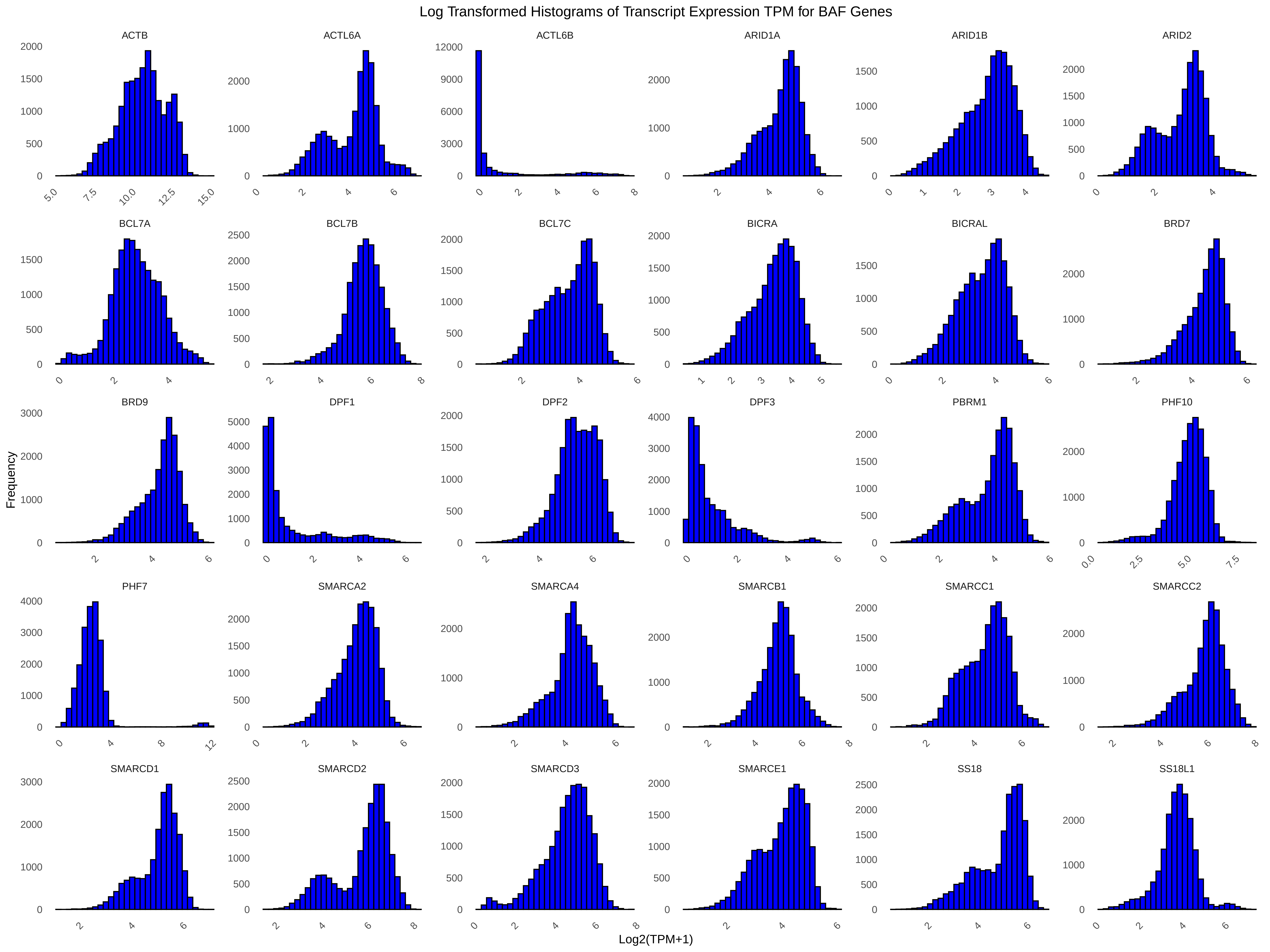

### Supplymemtal Figure S3

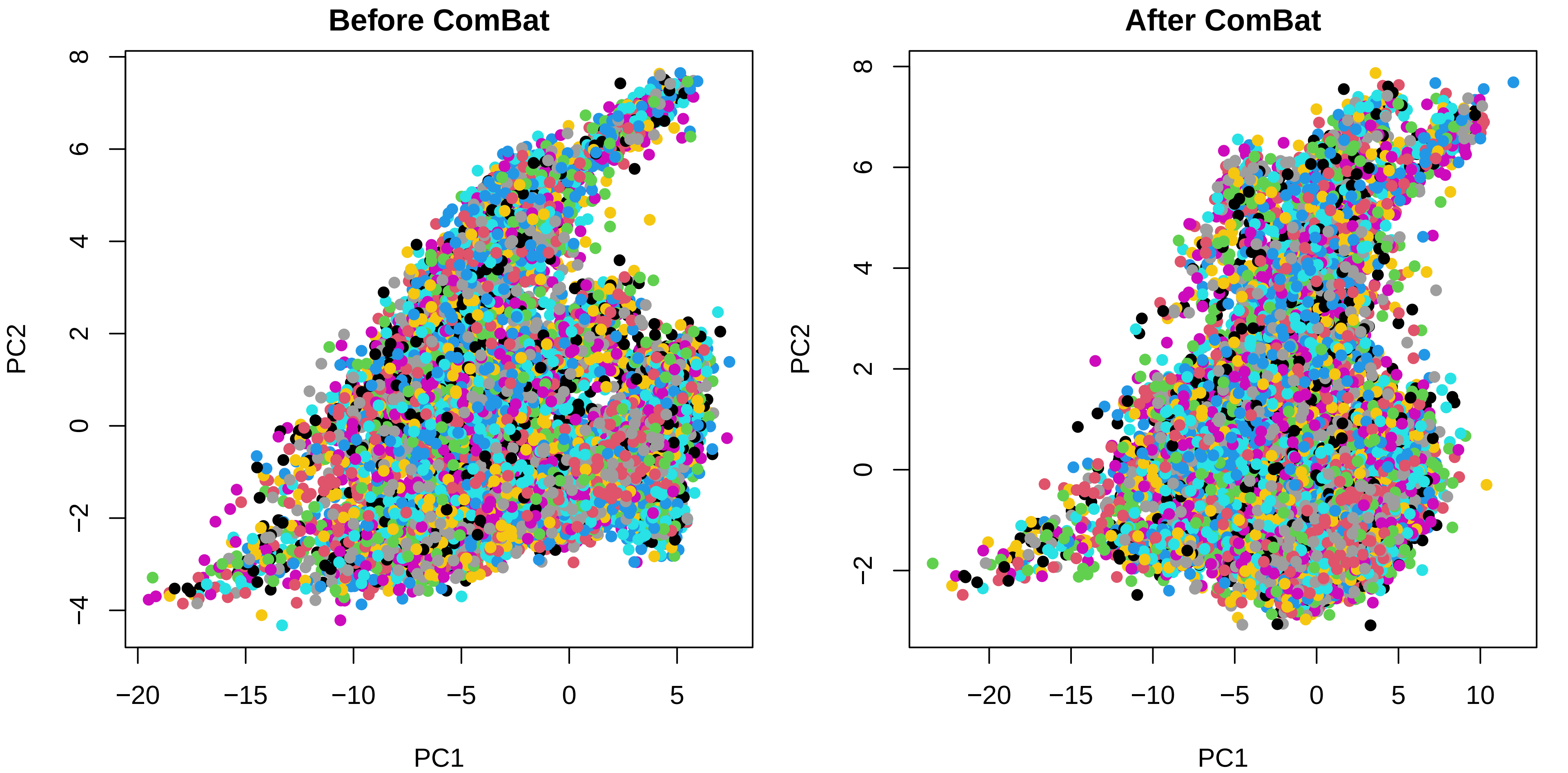

### Supplymemtal Figure S4

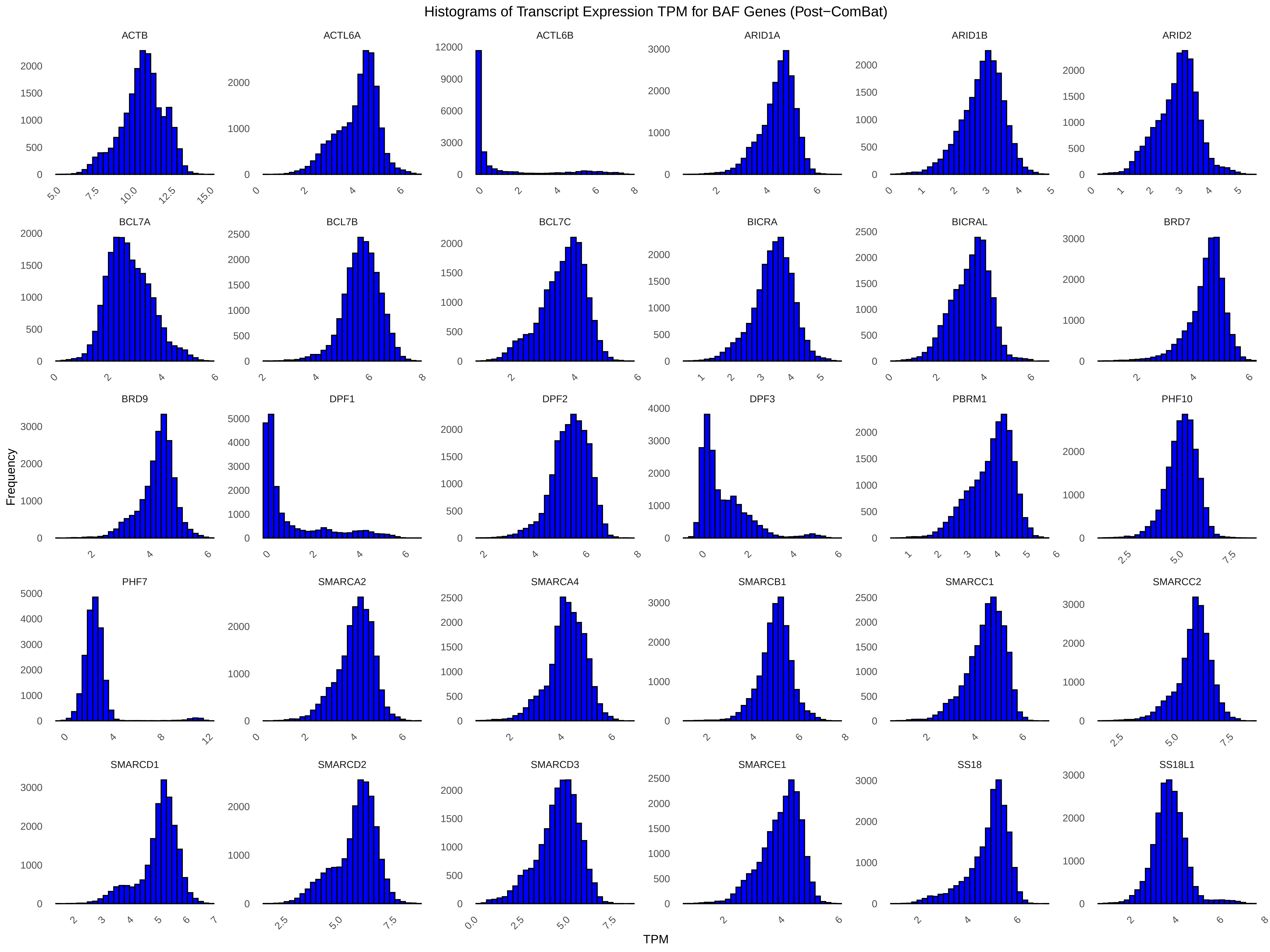

### Supplymemtal Figure S5

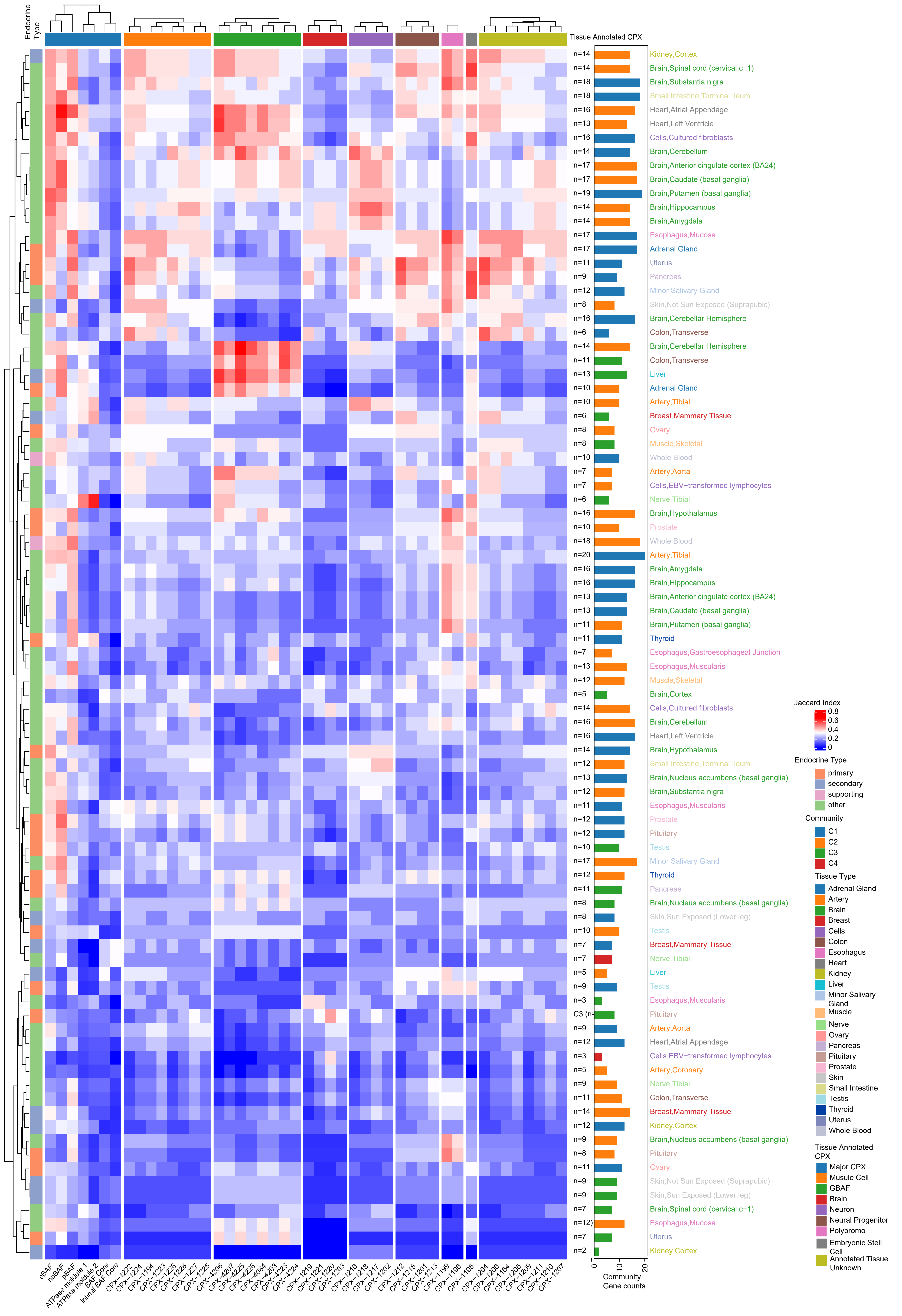
