## Supplementary material for "Tissue-Specific Co-Expression Patterns of BAF Complexes Provide Regulatory Insights Across Human Tissues with Implications for Endocrine and Non-Endocrine Functions": Supplymentary files explaination

**Supplementary Materials**

**Additional Figures S1–S4. Quality Assessment of BAF Gene Expression Data**

**Additional Figures S1.** Histograms of raw TPM values for the 30 BAF complex genes across all GTEx samples, illustrating the skewed expression distributions.

**Additional Figure S2.** Histograms of $\log_2(TPM + 1)$–transformed expression for the same genes, showing variance stabilization and reduced skew.

**Additional Figure S3.** Principal component analysis of $\log_2$-transformed BAF gene expression before (left) and after (right) batch-effect correction, demonstrating removal of sequencing-center effects.

**Additional Figure S4.** Histograms of $\log_2(TPM + 1)$ expression following batch-effect correction, confirming preservation of stabilized distributions for downstream network analysis.

**Additional Figure S5.** serves as the supplementary data source for **Figure 5**, displaying a heatmap of Jaccard similarity between GTEx tissue-based BAF gene communities (rows) and curated CPX reference subcomplexes (columns), which are grouped by subcomplex category (e.g., Major CPX, Muscle cell, GBAF, Brain, Neuron, Neuronal Progenitor, Polybromo-associated, Embryonic). Each row represents one of 49 GTEx tissues, annotated with endocrine classification and gene counts. Cell color reflects the Jaccard index, indicating the degree of overlap between each tissue's gene module and each reference complex, and enabling identification of major tissue clusters (**Blocks 1a, 1b, 1c, 2, and 3**). Due to figure size and resolution limitations, the CPX ID column is not shown in Figure 5. However, **Additional Figure S5** presents the same data structure and layout and includes the CPX column to explicitly indicate the reference BAF complex (CPX ID) for each comparison.

**Additional Table S1**: Final List of reference BAF protein complexes genes annotated by Tissue

We evaluated the concordance between identified gene communities and 46 curated reference BAF protein complexes across 45 tissues (see **Figure 3 and Additional Table S1**). Of the 46 reference complexes, the major types—including cBAF, ncBAF, pBAF, BAF cores, initial BAF cores, ATPase module 1, and ATPase module 2—were sourced from Mashtalir et al. [2], while the remaining 39 were obtained from the Complex Portal database .**Additional Table S1** lists the reference BAF complex (CPX) annotations, the associated tissues, gene members, and CPX IDs used for the concordance analysis. **Figure 3** provides a visual representation of these results. Of the 46 reference BAF protein complexes, the major complexes—including cBAF, ncBAF, pBAF,

BAF cores, initial BAF cores, ATPase module 1, and ATPase module 2—were sourced from Mashtalir et al., while the remaining 39 reference complexes were obtained from the Complex Portal database.

Complex Portal - CPX-1203

Complex Portal - CPX-1204

Complex Portal - CPX-1205

Complex Portal - CPX-1206

Complex Portal - CPX-1207

Complex Portal - CPX-1209

Mashtalir N, D'Avino AR, Michel BC, et al. Modular Organization and Assembly of SWI/SNF Family Chromatin Remodeling Complexes. Cell 2018 175 1272–1288.e20

**Additional Table S2:** Comprehensive Jaccard Index and P-Value Matrix for All Tissue–BAF Complex Associations (45 Tissues × 46 Complexes). **Tables 1.1–1.5** present only those tissue–complex pairs from **Additional Table S2** that met the criteria of Jaccard Index > 0.4 or P-Value Jaccard < 0.05. A full comparison across all 45 tissue types and 46 reference BAF complexes is available in **Additional Table S2** in the Supplementary Document.
